# Galectin-9 signaling drives breast cancer invasion through matrix

**DOI:** 10.1101/2021.11.06.467574

**Authors:** Dharma Pally, Mallar Banerjee, Shahid Hussain, Rekha V Kumar, Alexandra Petersson, Ebba Rosendal, Ludvig Gunnarsson, Kristoffer Peterson, Hakon Leffler, Ulf J. Nilsson, Ramray Bhat

## Abstract

Aberration in expression and function of glycans and their binding proteins (lectins) in transformed cells constitutes one of the earliest discovered hallmarks of cancer. Galectins are a conserved family of lectins that can bind to β-galactosides. Among them, the role of Galectin-9, a galectin with two carbohydrate binding domains in immune-tumor cell interactions has been well-established, although its effect on cancer cell behavior remains as yet unclear. In this study, we used a spectrum of cell lines from homeostatic breast cells to transformed non-invasive and invasive cell lines cultured in microenvironment-diverse conditions to show that Galectin-9 expression shows an elevation in association with invasiveness of breast cancer epithelia. Our observations were supported by immunohistochemical studies of breast tumors and adjacent normal-tissues from patients. Genetic perturbation of Galectin-9 as well as the pharmacological inhibition of activity using cognate inhibitors confirmed a positive correlation between Galectin-9 levels and the adhesion of the aggressive triple negative breast cancer cells MDA-MB-231 to- and their invasion through-extracellular matrices (ECM). Within a constituted organomimetic multiECM microenvironment, Galectin-9 enhanced both the solitary and the collective invasion of cancer cells. Quantitative proteomics led us to uncover the inductive role of Galectin-9 in the expression of the proinvasive protein S100A4. In addition, Galectin-9 expression correlated with FAK signaling, the inhibition of which decreased S100A4 mRNA levels. Our results provide crucial signaling insights into how the elevation in Galectin-9 expression in breast cancer cells potentiates their invasiveness through ECM during early steps of metastasis.

## Introduction

Galectins are a family of soluble lectins with a characteristic (about 135 amino acid) carbohydrate-binding domain (CRD) that binds to β-galactoside-containing glycans (1,2). Widely expressed across tissues, galectins are seen both inside cells and in the extracellular space, and based on their location, modulate a diverse set of functions including, but not limited to, cell proliferation, apoptosis, pre-mRNA splicing, cell-cell and cell-ECM (extracellular matrix) adhesion, epithelial cell polarity, innate/adaptive immunity regulation, and migration (3-7). Given that these phenomena contribute to tissue architecture and homeostasis, aberrant expression of galectins have been increasingly found to be associated with carcinogenesis. In fact, galectins have been shown to regulate tumor cell growth, apoptosis, invasion, and metastasis, oncogenic signaling and immune suppression (3,8,9).

Galectin-9 belongs to a subset of the 15 mammalian galectins, with two different CRDs in one peptide chain, here termed N-CRD and C-CRD (10). Rigorous investigations on its functions have predominantly been confined to immunological contexts, such as the activation of dendritic cells and their induction of naïve T cell proliferation, a level dependent modulation of activated T cell fate and the regulation of B-cell driven autoimmunity (11-13).

The role of Galectin-9 in breast cancer progression is as yet unclear. Although early studies showed an association of its expression with better prognosis in breast cancer patients, an increasing set of investigations implicate Galectin-9 as an invasion-promoting protein through its role in mediating the escape of cancer cells from immune surveillance. Recent evidence shows that the expression of Galectin-9 in cancers of breast, colon and in myeloid leukemia is regulated by proinvasive microenvironmental cues such as HIF-1α and TGFβ (14). Overexpression of Galectin-9 in breast cancer has also been reported by Grosset and coworkers, where the subcellular localization of the protein was found to affect the prognosis of patients (15). In the context of the well-known demonstration of localization-specific roles of other galectins, and isoform-diverse expression of Galectin-9, a careful investigation of the effect of the latter on transformed cell phenotype is clearly warranted (15-17).

In this paper, we probed the expression of Galectin-9 within sections of patients with different breast cancer histological subtypes as well as an invasion-diverse set of breast cancer cell lines cultured on organotypic extracellular matrix (ECM) scaffolds. We show that the levels of Galectin-9 correlate with the ability of breast cancer epithelia to adhere to ECM as well as invade both collectively and as dispersed mesenchymal cells within reconstructed organotypic ECM-complex microenvironments. The potentiation of invasiveness is dependent on its upregulation of FAK signaling, which in turn regulates the metastasis-associated protein S100A4.

## Results

### Galectin-9 is highly expressed in breast cancer epithelia

We began our investigation by assaying Galectin-9 expression across four breast epithelial cell lines: a) an immortalized human mammary epithelial cell line (HMLE) (typifying untransformed breast epithelial cells, given its ability to form growth-arrested lumen-containing acini when cultured in mammotypic laminin-rich ECM), b) MCF7 cells (typifying non-invasive tumor cells that form a dysmorphic solid cluster in lrECM), and c) BT-549 and MDA-MB-231 cells (which represent full blown stromal invasive cancer cells that form stellate invasive morphologies within lrECM).

Galectin-9 gene expression was measured in the cells cultured as monolayers (Figure 1A) and on two ECM scaffolds using qRT-PCR: the nonfibrillar lrECM to mimic the milieu of cells in homeostasis or just before the basement membrane (BM) is breached (Figure 1B) and the fibrillar Collagen I (Coll I), the stromal ECM milieu through which cancer cells invade (Figure 1C). Galectin-9 transcript levels were significantly higher in MDA-MB-231 cultured on plastic, lrECM and Coll I and in BT-549 cultured on plastic and Coll I compared with MCF7 and HMLE cells, which showed relatively lower levels of Galectin-9 mRNA (Figure 1A-C and Figure S1A). The upregulation of Galectin-9 mRNA in MDA-MB-231 and BT-549 was consistent with higher levels of proteins in these cells as observed through immunoblotting, whereas HMLE and MCF7 showed sparse signals (Figure 1D). To investigate if Galectin-9 levels were elevated in breast tumors *in vivo*, we stained histological sections from a cohort of 11 breast cancer patients along with their matched non-malignant tissues. Primary antibody-omitted controls were used to confirm the veracity of staining (Figure S1B). Out of 11, 10 samples showed increased Galectin-9 levels in tumors compared with matched adjacent normal tissues (across all histotypes except in HER2^+^ subtype where Galectin-9 staining was elevated in tumors, but matched non-cancerous tissue was not available) (Figure 1E-G & S1C). Our results were further confirmed by observation of an increased Galectin-9 mRNA and protein levels within breast tumors, when examined within publicly available samples curated by The Cancer Genome Consortium (TCGA; using RNA sequencing) and the Clinical Proteomic Tumor Analysis Consortium (CPTAC using a combination of large-scale proteome and genome analysis) respectively (Figure S1D and E). In fact, its mRNA levels were found to be higher in breast cancer samples from stages 1,2 and 3 compared with adjacent non-cancerous control samples (Figure S1F). Among the histopathological types, the most prevalent type: invasive ductal carcinoma (IDC) samples showed widely variable but significantly higher Galectin-9 mRNA and protein levels compared with noncancerous controls within the above-mentioned databases (Figure S1G & S1H). Interestingly, within the TCGA dataset, we observed a significant hypomethylation of *LGALS9* promoter within primary tumor samples compared with controls, suggesting a possible epigenetic upregulation of Galectin-9 associated with breast cancer progression (18,19) (Figure S1I).

**Figure 1:**
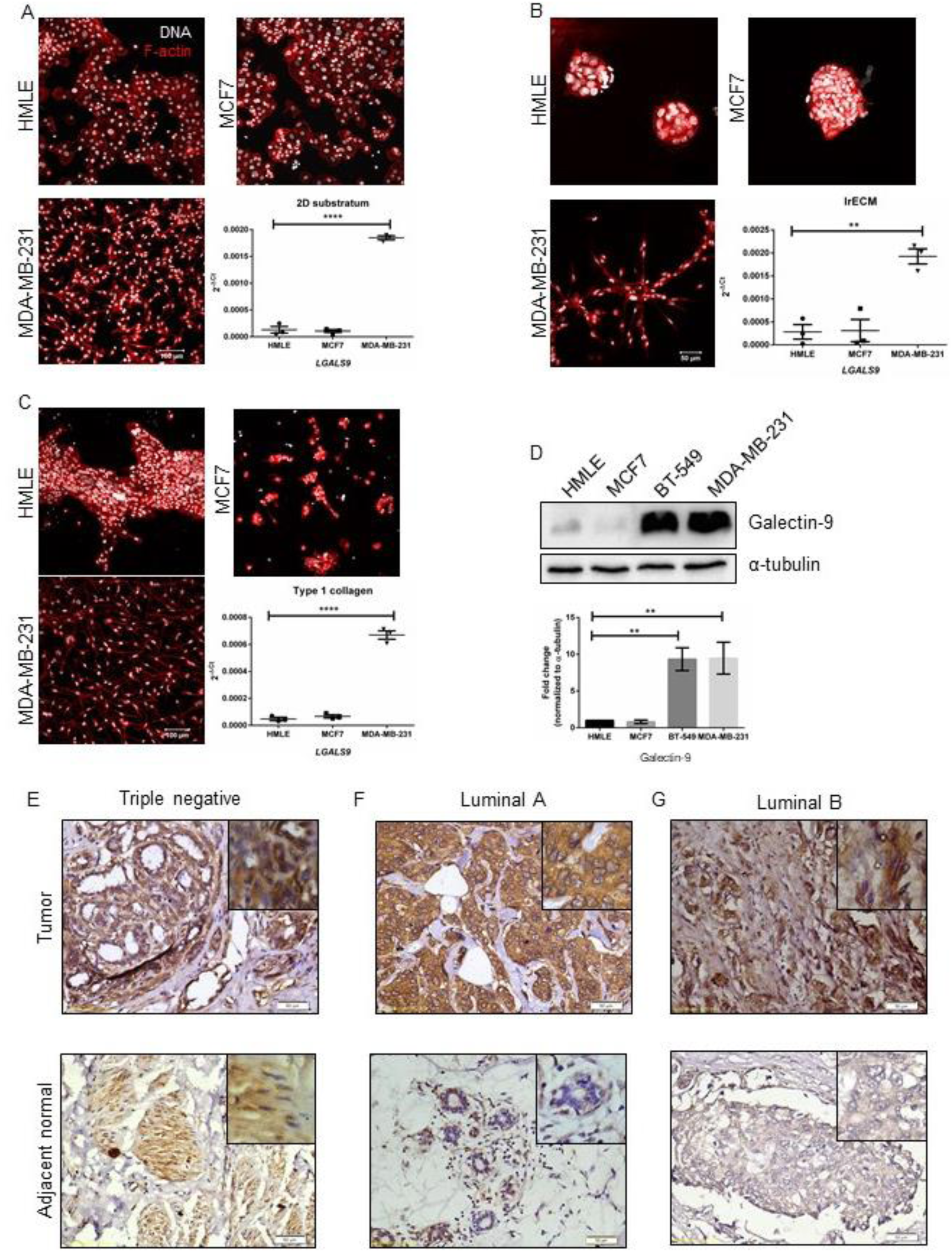
Galectin-9 expression positively correlates with the invasiveness of breast cancer cells. A) Representative micrographs showing characteristic cell morphology of HMLE (top, left), MCF7 (top, right), and MDA-MB-231 (bottom, left) when grown on 2D substratum. Scale bar: 100 μm. MDA-MB-231 cells show significantly higher Galectin-9 gene expression compared to HMLE cells when cultured as monolayer (bottom, right). B) Representative micrographs showing characteristic cell morphology of HMLE (top, left), MCF7 (top, right), and MDA-MB-231 (bottom, left) when grown on nonfibrillar lrECM scaffold. Scale bar: 50 μm. Bar graph showing significantly higher Galectin-9 transcript levels in MDA-MB-231 compared to HMLE cells when cultured on nonfibrillar lrECM scaffold (bottom, right). C) Representative micrographs showing characteristic cell morphology of HMLE (top, left), MCF7 (top, right), and MDA-MB-231 (bottom, left) when grown on fibrillar Collagen I scaffold. Scale bar: 100 μm. Bar graph showing significantly higher Galectin-9 transcript levels in MDA-MB-231 compared to HMLE cells when cultured on fibrillar Collagen I scaffold (bottom, right). *18S rRNA* is used as internal control to normalize Galectin-9 transcript levels. n=3, Mean ± SEM. **P≤0.01, ****P<0.0001. D) Immunoblot showing higher Galectin-9 levels in BT-549 and MDA-MB-231 compared to HMLE and MCF7 cells (top). Bar graph showing quantification of Galectin-9 levels in HMLE, MCF7, BT-549 and MDA-MB-231. Normalized to α-tubulin. n=3, Mean ± SEM. **P≤0.01 (E-G) Representative immunohistochemistry images of Galectin-9 staining in breast cancer tissue (top) and adjacent normal (bottom) in triple negative, Luminal A and Luminal B subtypes respectively. Galectin-9 levels are elevated in tumor tissues of all the subtypes of breast cancer compared to matched normal tissues. Scale bar: 50 μm. One-way ANOVA with Dunnett’s multiple comparisons.

### Decrease in Galectin-9 expression and function decreases adhesion of breast cancer cells to ECM and their invasion through pathotypic ECM microenvironments

Having established that Galectin-9 levels were elevated in breast cancer cells, we sought to stably knock down its expression in MDA-MB-231 cells using a lentiviral transduction of gene-cognate shRNA (scrambled non-specific shRNA denoted in figures as shSc was used as control). Knockdown was confirmed using qRT-PCR and immunoblot (Figure S2A & S2B). Galectin-9-depleted cancer cells were observed to invade to a lower extent through lrECM-coated transwells, when compared with control cells (Figure 2A). In an earlier work, we have observed that invasion through ECM for MDA-MB-231 cells directly correlates with its ability to adhere to ECM substrata (20). Therefore, we assayed for, and observed that, the adhesion of cells to lrECM- and Coll I-coated substrata was significantly reduced upon depletion of Galectin-9 (Figure 2B & 2C).

**Figure 2:**
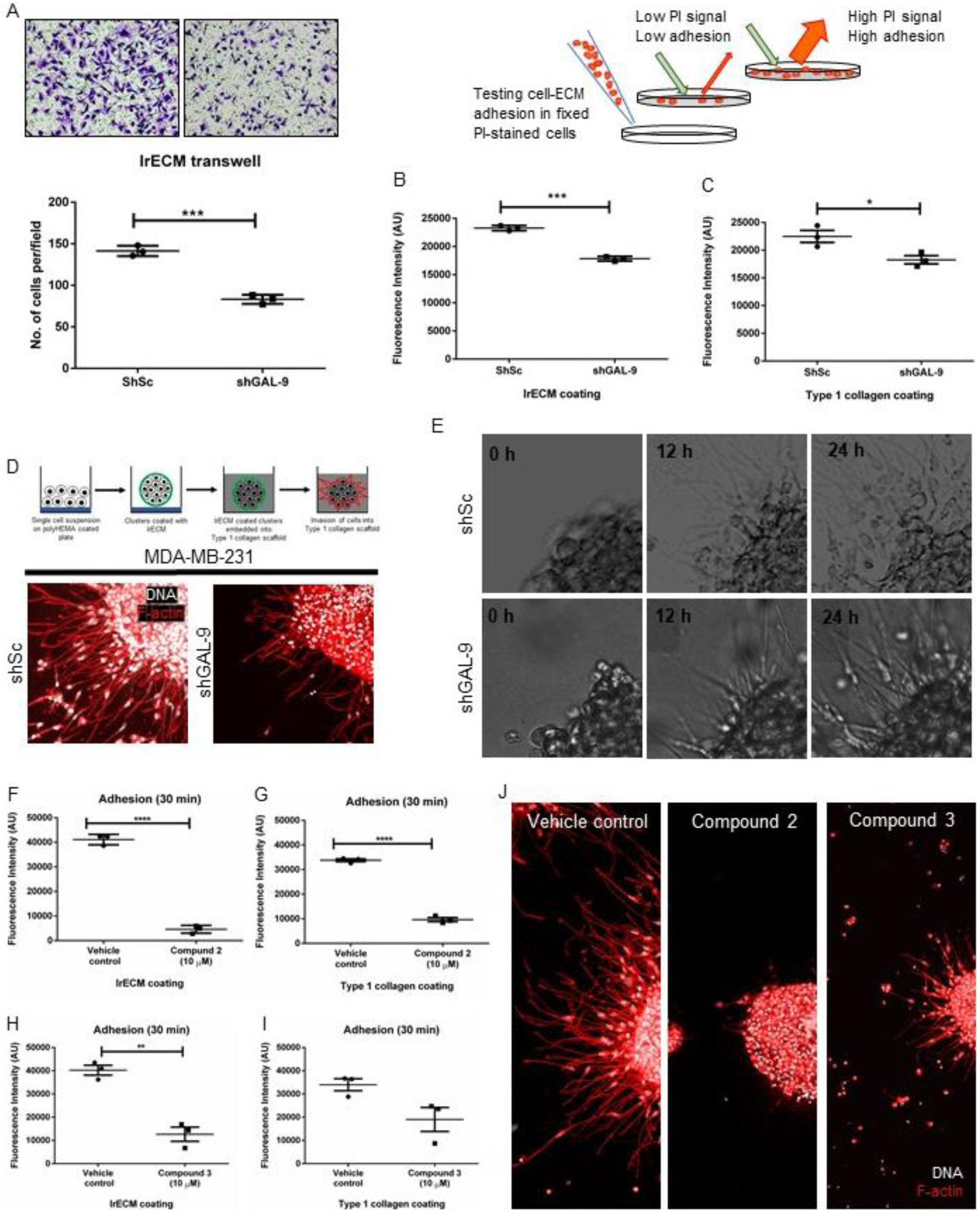
Decreased breast cancer invasion upon genetic perturbation and functional inhibition of Galectin-9. A) Graph showing a significant decrease in invasion upon Galectin-9 knockdown compared to scrambled control cells through lrECM coated transwells. B) Graph depicting adhesion of MDA-MB-231 cells to lrECM. Cells upon Galectin-9 knockdown show a significant decrease in adhesion to lrECM. C) Graph depicting adhesion of MDA-MB-231 cells to Coll I. Cells upon Galectin-9 knockdown show a significant decrease in adhesion to Coll I. D) Maximum intensity projection of confocal micrograph showing decreased invasion of cancer cells after Galectin-9 knockdown (right) compared to scrambled control cells (left). Cells are counterstained for nucleus (grey) with DAPI and F-actin (red) with Phalloidin. Scale bar: 200 μm. Schematic adapted from (23) E) Bright-field images taken at 0 h, 12 h, and 24 h from time-lapse videography of lrECM-coated clusters of scrambled control (top) and Galectin-9 knockdown cells (bottom) invading into surrounding Coll I. Scale bar: 50 μm. F) Graph showing significant decrease in breast cancer cells adhesion to lrECM after inhibiting Galectin-9 using **2** (10μM). G) Graph showing significant decrease in breast cancer cells adhesion to Coll I after inhibiting Galectin-9 using **2** (10μM). H) Graph showing significant decrease in breast cancer cells adhesion to lrECM after inhibiting Galectin-9 using **3** (10μM). I) Graph showing decrease in breast cancer cells adhesion to Coll I after inhibiting Galectin-9 using **3** (10μM). J) Maximum intensity projection of confocal micrograph showing decreased invasion of cancer cells after Galectin-9 inhibition using **2** and **3** compared to vehicle treated cells. n=3, mean ± SEM, * P≤0.05, ** P≤0.01, ***P≤0.001, **** P<0.0001. Unpaired Student’s *t*-test with Welch’s correction.

Invasive breast cancer epithelia are capable of migrating within ECM using a combination of distinct modes: collective and mesenchymal single-cell invasion (21,22). This is evident when the cells are topologically surrounded by an outwardly radial ECM arrangement of BM and then Coll I, similar to what is seen in vivo (23). Accordingly, cancer cell clusters were prepared using Galectin-9-depleted and control cells, and after coating with lrECM, embedded in Coll I scaffolds as described earlier (see also Figure 2D top). It was evident from maximum intensity projections of confocal micrographs taken after 24 hours, that Galectin-9 knockdown significantly impaired cancer cell invasion into Coll I scaffolds (Figure 2D bottom right) compared to control cells (Figure 2D bottom left). A time-lapse video microscopic examination also revealed slower and sparser egress of breast cancer cells from the cluster into the surrounding collagenous microenvironment (Figure 2E).

We sought to validate our findings in the context of the knockdown by designing and evaluating Galectin-9 specific inhibitors that impair its binding to β-galactosides. The design of novel more selective and potent Galectin-9 inhibitors was inspired by the recent publications that halophenyl thio-α-D-galactopyranosides show significant affinity enhancement over simple galactosides for most galectins (24) and that galactosides carrying a *N*-sulfonylamidine moiety at O3 display increased affinity and high selectivity for the N-terminal domain of Galectin-9 (25). Synthesis of two novel 3,4-dichlorophenyl thio-α-D-galactopyranosides carrying different N-sulfonylamidine moieties at O3, **2** and **3**, were performed as reported (25) from the known 3,4-dichlorophenyl 3-*O*-propargyl-thio-α-D-galactopyranoside **1** (Scheme 1) (26). Evaluation of galectin affinities were performed as reported (25,27) and clearly revealed both compounds being selective inhibitors of galectin-9 N-CRD with low μM affinities (Table 1). Compound **2** is somewhat more potent than **3**, while both show good, but to some extent different selectivity for Galectin-9 N-CRD over other galectins. Hence, **2** and **3** are suitable small-molecule inhibitors for evaluating effects of pharmacological inhibition of Galectin-9 in cell assays.

**Scheme 1.**
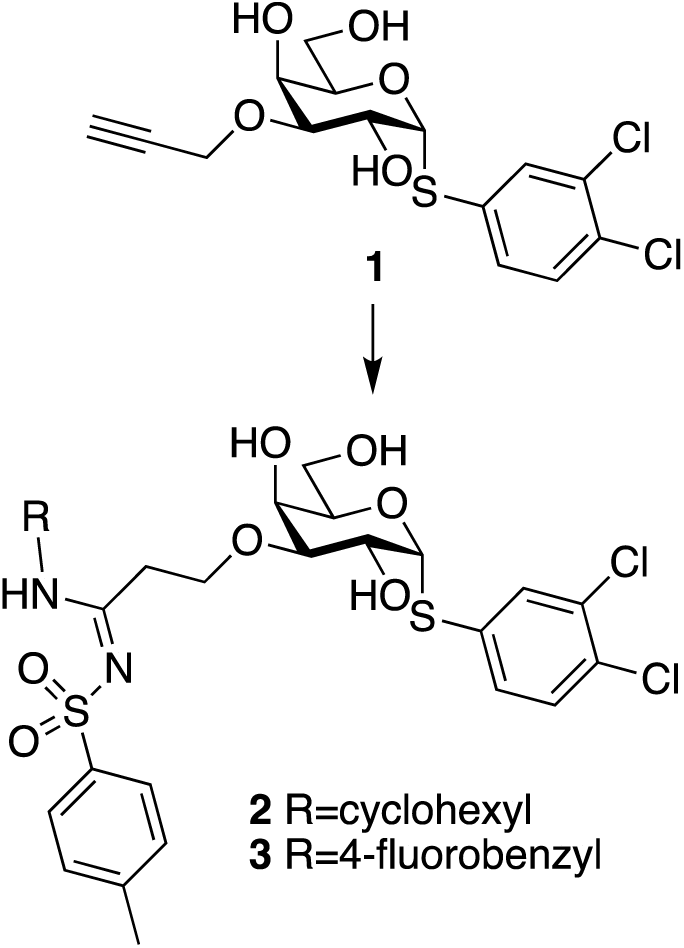
Synthesis of selective Galectin-9N inhibitory sulfonylamidine-appended galactosides 2 and 3. Reaction conditions: THF, H_2_O, tosyl azide, cyclohexylamine or 4-fluorobenzylamine, CuI, 100°C, 50% of **2**, 6.7% of **3**.

**Table 1:**
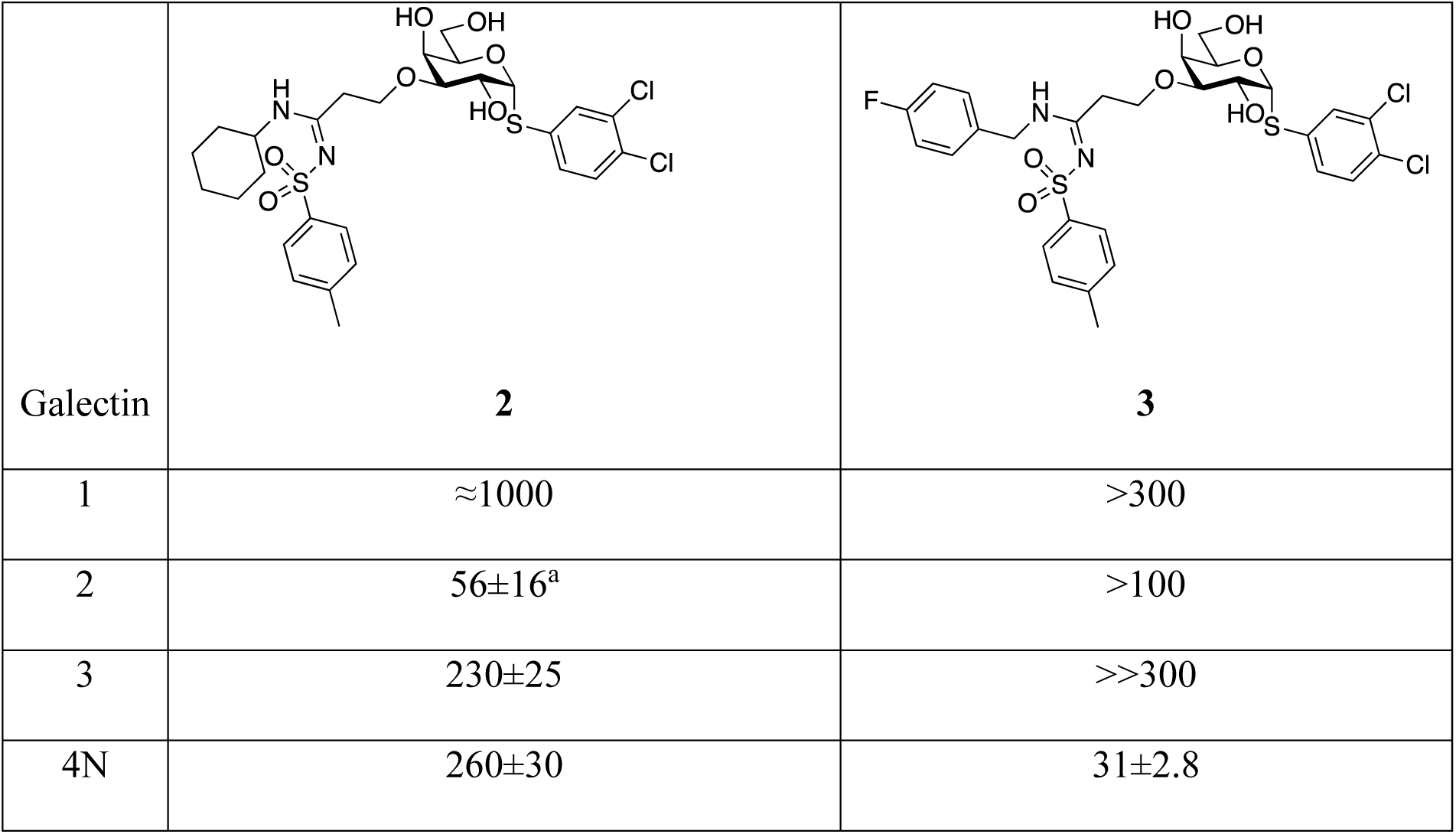
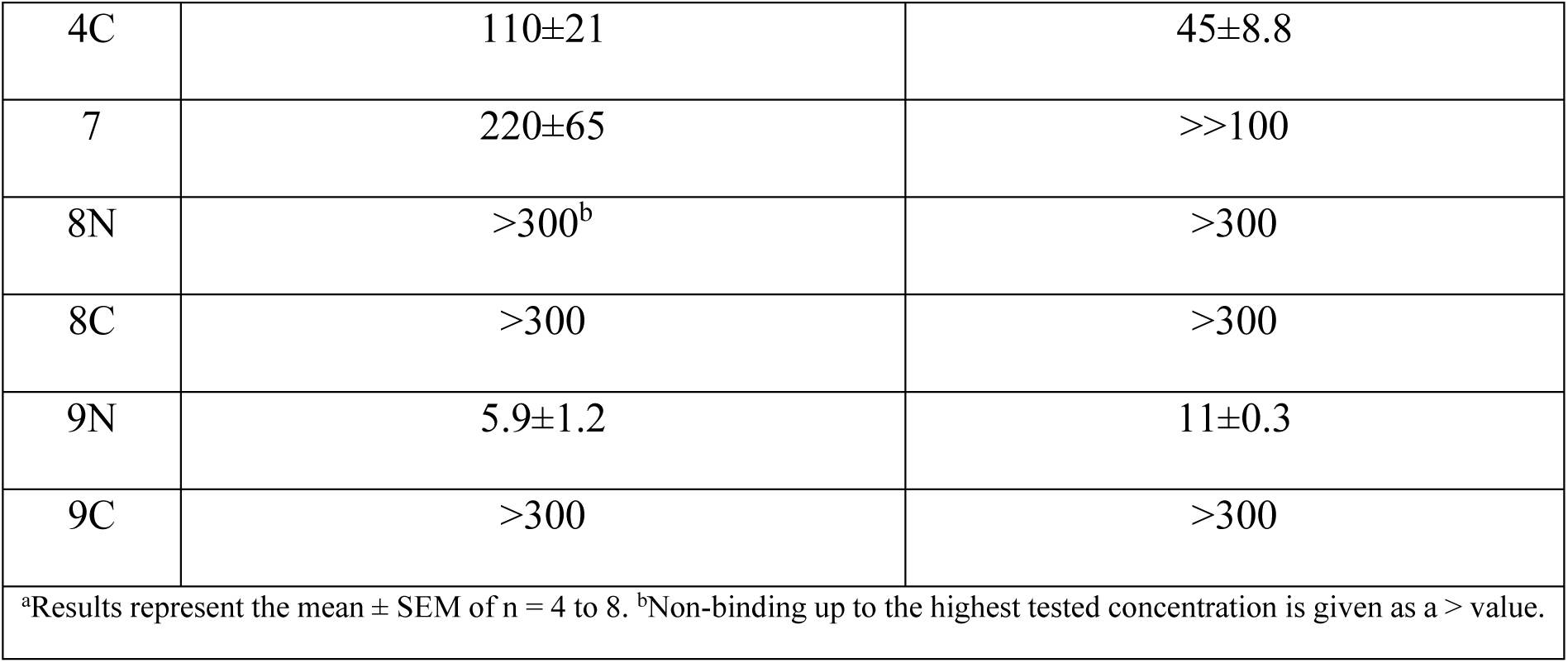
Structures of **2** and **3** and dissociation constants (K_d_ in μM) against human galectin-1, 2, 3, 4N (N-terminal domain), 4C (C-terminal domain), 7, 8N, 8C, 9N, and 9C

| Galectin | **2** | **3** |
| --- | --- | --- |
| 1 | $\approx 1000$ | $>300$ |
| 2 | $56 \pm 16^a$ | $>100$ |
| 3 | $230 \pm 25$ | $>>300$ |
| 4N | $260 \pm 30$ | $31 \pm 2.8$ |
| 4C | 110±21 | 45±8.8 |
| 7 | 220±65 | >>100 |
| 8N | >300<sup>b</sup> | >300 |
| 8C | >300 | >300 |
| 9N | 5.9±1.2 | 11±0.3 |
| 9C | >300 | >300 |
| <sup>a</sup>Results represent the mean ± SEM of n = 4 to 8. <sup>b</sup>Non-binding up to the highest tested concentration is given as a > value. |  |  |

Toxicity of compounds **2** and **3** on MDA-MB-231 cells was assessed after 24 hours treatment and IC50 values were determined to be close to the K_d_ values determined in the protein binding fluorescence polarization assay: 18 μM & 19 μM for **2** and **3**, respectively (Fig S2C and S2D). We treated MDA-MB-231 cells with a sub-IC50 concentration of **2 & 3** (10 μM) for 1 hour prior to performing adhesion on lrECM and Coll I. Inhibition of Galectin-9 using either of the two inhibitors significantly decreased cancer cell adhesion to both lrECM and Coll I matrix when compared with vehicle-treated cells (Figure 2F-I). Subsequently, we assayed for cancer epithelial invasion through 3D multi matrix scaffolds in the presence of Galectin-9 inhibition. Similar with Galectin-9 knockdown, inhibition with **2** and **3** abrogated cancer cell migration into Coll I compared with vehicle-treated cells (Figure 2J). Our genetic perturbation and small molecule inhibitor experiments suggest that Galectin-9 might play an inductive role in early cancer cell invasion.

### Galectin-9 overexpression increases cancer cell adhesion and invasion through matrix

To understand if increased Galectin-9 positively correlates with invasive potential of cancer cells, we overexpressed its cDNA in MDA-MB-231 cells and confirmed its increased expression (Fig S3). Overexpression of Galectin-9 led to an increase in the invasion of MDA-MB-231 cells through lrECM-coated transwells (Figure 3A). Next, we assayed for adhesion of cancer cells to lrECM and Coll I. Overexpression resulted in increased adhesion of cancer epithelia to both the extracellular matrices (Figure 3B & 3C). Invasion through transwells does not address the mode of cancer cell invasion that is altered. To understand this, we embedded cancer cell clusters coated with lrECM in polymerizing Coll I. At the end of 24 hours, Galectin-9 overexpressing cells showed higher invasion into Coll I when compared to empty vector overexpressing cells (Figure 3D). With the help of time-lapse microscopy, we quantified collective cell invasion and single cell invasion of cancer epithelia (Figure 3E). Galectin-9 overexpression significantly increased both collective and single cell modes of invasion when compared to the control cells (Figure 3F & 3G). Galectin-9 also regulated the velocity of single cells migrating in Coll I leading to increased accumulated distance of cells (Figure 3H & 3I).

**Figure 3:**
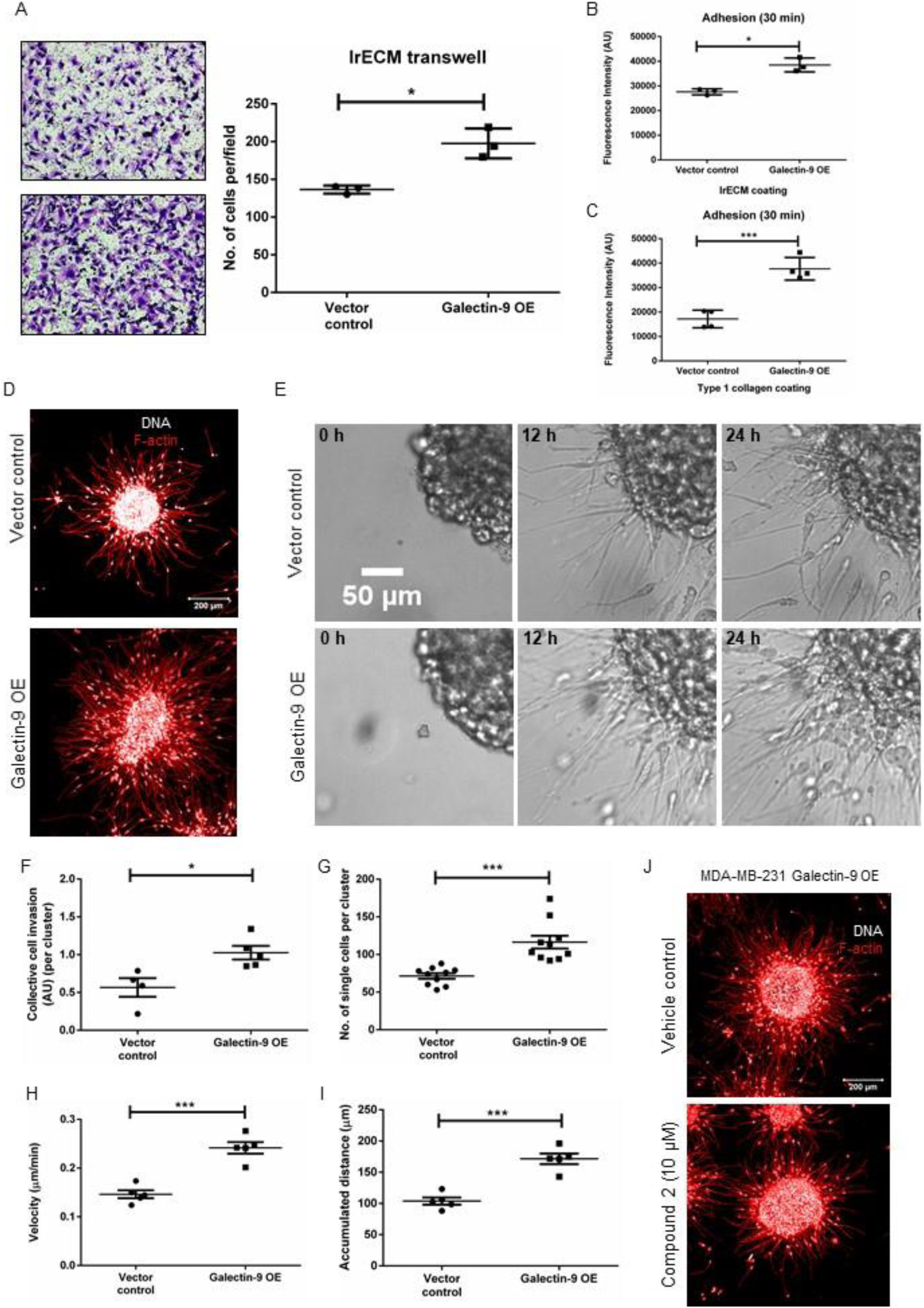
Increased Galectin-9 expression concomitantly increased both collective and single-cell invasion. A) Graph showing a significant increase in invasion of Galectin-9 overexpressing cells compared to vector control cells through lrECM coated transwells. B) Graph depicting adhesion of MDA-MB-231 cells to lrECM. Galectin-9 overexpression significantly increases the adhesion of cancer cells to lrECM. C) Graph depicting adhesion of MDA-MB-231 cells to Coll I. Galectin-9 overexpression significantly increases the adhesion of cancer cells to the matrix. n≥3, Mean ± SEM. D) Maximum intensity projection of confocal micrograph showing higher invasion of MDA-MB-231 cells overexpressing Galectin-9 (bottom) into Coll I compared to vector control cells (top). Cells are counterstained for nucleus (grey) with DAPI and F-actin (red) with Phalloidin. Scale bar: 200 μm. E) Bright-field images taken at 0 h, 12 h, and 24 h from time-lapse videography of lrECM-coated clusters of vector control and Galectin-9 OE (top and bottom, respectively) invading into surrounding Coll I. Scale bar: 50 μm. F) Graph depicting significant differences in collective cell mode of invasion of vector control and Galectin-9 OE cells as measured by increased cluster size obtained from time-lapse videography. G) Graph depicting significant differences in single-cell mode of invasion of vector control and Galectin-9 OE cells as measured by number of dispersed single cells in Coll I. H) Graph showing significantly higher mean migratory velocity of single Galectin-9 OE cells than vector control cells as measured by manual tracking dynamics obtained from lapse videography. I) Graph showing significantly higher mean accumulated distance of single Galectin-9 OE cells than vector control cells as measured by manual tracking dynamics obtained from lapse videography. n≥3, N=50, mean ± SEM. J) Maximum intensity projection of confocal micrograph of Galectin-9 overexpressing cells showing decreased invasion after Galectin-9 inhibition using **2**. * P≤0.05, ***P≤0.001. Unpaired Student’s *t*-test with Welch’s correction.

Next, we asked if the increased cancer cell invasion shown by Galectin-9 overexpressing cells could be reversed by a cognate pharmacological impairment? To address this, we used the inhibitor **3** and allowed cancer cells to invade through multi-ECM scaffolds. Inhibition of elevated Galectin-9 levels decreased cancer epithelial invasion when compared to vehicle-treated Galectin-9 overexpressing cells (Figure 3J). The reduced invasion upon inhibitor treatment was comparable to the vector control cells in Figure 3D. Our observations thus confirmed that Galectin-9 levels positively correlated with the invasive potential of cancer cells and might play an inductive role during breast cancer cell invasion. In addition, the pharmacological treatments suggested that such a correlation was dependent on the carbohydrate-binding activity of Galectin-9.

### Galectin-9 potentiates cancer cell invasion by up-regulating S100A4 and focal adhesion kinase activation

We next sought to understand how Galectin-9 regulated cell adhesion to lrECM and Coll I, and invasion through multi-ECM environments. A quantitative proteomics approach was taken to identify the proteins that are dysregulated when Galectin-9 was overexpressed. We observed that 123 proteins were differentially expressed to significant extents (either upregulated or downregulated by 1.5-fold) (Figure 4A). Functional annotation using ontological analysis showed such proteins to be significantly involved in cell-cell adhesion, cytoskeleton organization, and intermediate filament organization (Figure 4B). Similarly, KEGG pathway analysis also predicted that the differentially regulated proteins are involved in tight junction, actin organization and transendothelial migration pathways (Figure 4C). We further probed proteins amongst top 20 upregulated proteins for validation using gene expression (Figure S4). Amongst the genes we analyzed, the expression of *S100A4*, a calcium binding protein was positively correlated with Galectin-9 levels (Figure 4D and 4E). In addition, inhibition of Galectin-9 using the inhibitors **2** and **3** resulted in significantly reduced S100A4 gene expression (Figure 4F).

**Figure 4:**
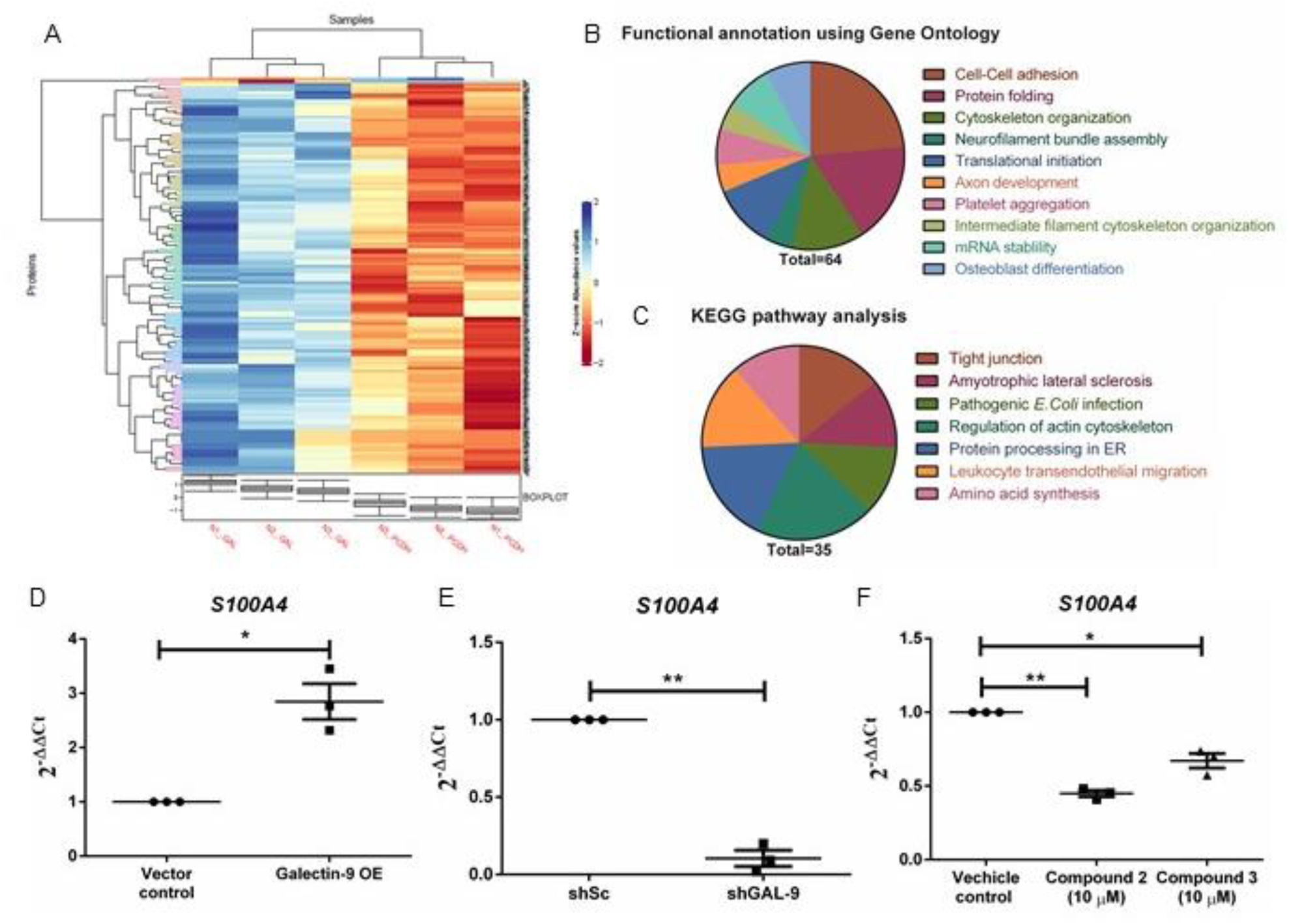
Galectin-9 mediates cancer invasion by up-regulating *S100A4*. A) Heat map showing list significantly up-regulated (blue) and down-regulated (red) proteins upon Galectin-9 overexpression using mass spectrometry. B) Pie chart showing functional annotation of the differentially expressed proteins using gene ontology. C) Pie chart showing the pathway annotation of the differentially expressed proteins using KEGG pathway analysis. D) Graph showing significantly increased *S100A4* expression upon Galectin-9 overexpression. E) Graph showing significantly decreased *S100A4* expression upon genetic knockdown of Galectin-9 expression. F) Graph showing significantly decreased *S100A4* expression upon functional inhibition of Galectin-9 using **2** (10 μM) and **3** (10 μM) inhibitors.

During cancer cell invasion, cell-ECM adhesion has been shown to play a prominent role in mediating mesenchymal mode of invasion (28,29). Integrins are one of the major players in mediating cell adhesion to ECM. Outside-in signaling through integrins allows the phosphorylation of focal adhesion kinase (FAK) which in turn regulates actin polymerization dynamics that play a crucial role in cancer cell invasion (28,30). Therefore, we sought to probe whether FAK activation could be modulated by Galectin-9 levels.

Upon perturbing Galectin-9 expression in MDA-MB-231 cells, we probed for pFAK (Y397) signals. Phosphorylation of FAK correlated with Galectin-9 levels i.e., over expression of Galectin-9 resulted in increased phosphorylation (Figure 5A). Similarly, when Galectin-9 levels were knocked down, phosphorylation of FAK decreased concomitantly (Figure 5B). Next, we probed for kinetics of pFAK upon inhibiting Galectin-9 using **3**. We did not observe any immediate changes in pFAK levels at an early time point (30 min); at later time points, 1 hour and 6 hour, pFAK levels reduced when compared to 0 hour (Figure 5C). These observations suggest that Galectin-9 regulates *S100A4* expression and phosphorylation of FAK that are instrumental to the modulation of cancer cell adhesion and invasion. We next sought to verify a potentially inductive effect of FAK signaling on S100A4 as observed in other studies (31,32): in Galectin-9 overexpressing cells, which were treated with a pharmacological inhibitor of FAK signaling (CAS 4506-66-5; 100 nM), we observed a moderate downregulation in *S100A4* mRNA compared with vehicle controls (Figure 5D). These observations suggest that the upregulation of the migration-potentiating *S100A4* by Galectin-9 is regulated by FAK signaling.

**Figure 5:**
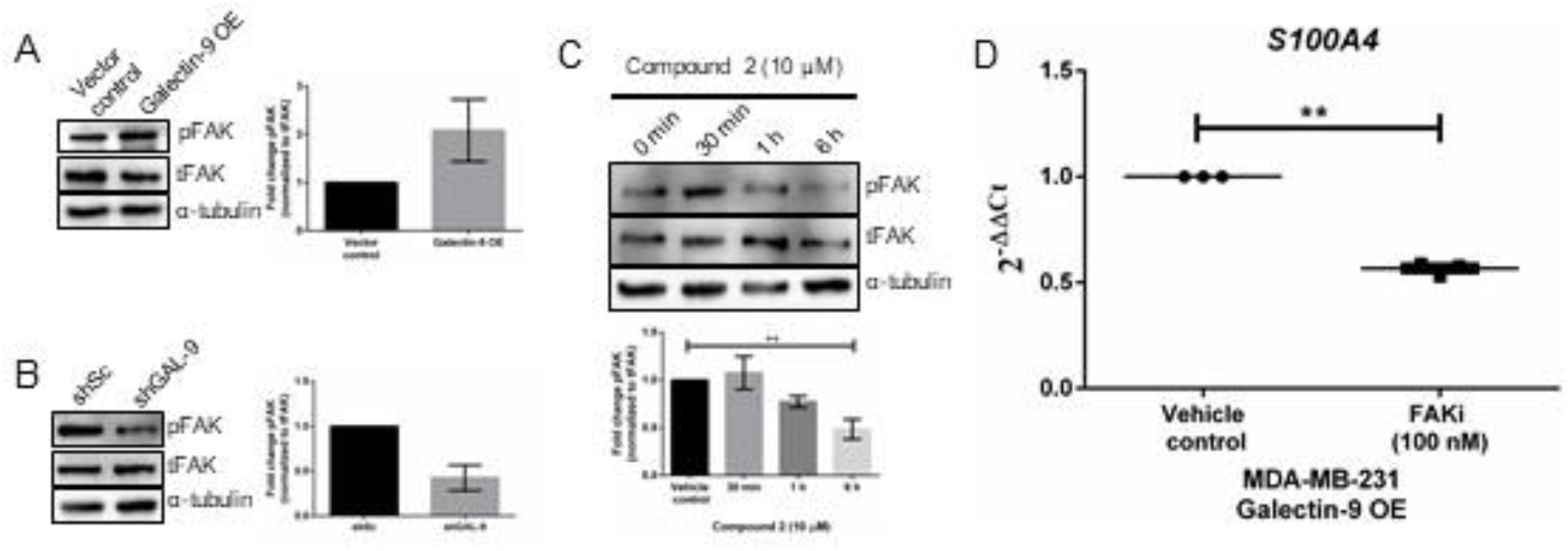
Galectin-9 regulates *S100A4* expression through FAK activation. A) Immunoblot showing increased phosphorylation of FAK upon Galectin-9 overexpression. Bar graph showing increased pFAK levels when Galectin-9 is overexpressed compared to vector control cells. B) Immunoblot showing decreased phosphorylation of FAK upon Galectin-9 knockdown. Bar graph showing decreased pFAK levels when Galectin-9 is knocked down compared to scrambled control cells. C) Immunoblot showing pFAK levels upon Galectin-9 functional inhibition using **2** (10 μM) at 0 h, 30 min, 1 h and 6 h. Bar graph showing decrease in pFAK levels at 1 hour and 6 h after inhibition of Galectin-9. D) Graph showing significant decrease in *S100A4* expression upon FAK inhibition (100 nM) compared to vehicle treated Galectin-9 OE cells. pFAK levels were normalized to total FAK. n=3, mean ± SEM, ** P≤0.01. One-way ANOVA with Dunnett’s multiple comparisons & unpaired Student’s *t*-test with Welch’s correction.

## Discussion

An earlier study on Galectin-9 in breast cancer observed that its levels positively correlate with cell aggregation, thereby decreasing dissemination (27). However, the cell line used in the study, MCF7 invades sparsely, if at all. In addition, RNAseq- and immunohistochemistry-based studies indicate an increase in Galectin-9 mRNA and protein levels at advanced stages of cancer progression (18). In the present study, we rigorously confirmed the increase in Galectin-9 both in breast cancer patient samples using immunohistochemistry as well as in an invasion-diverse set of established cancer (and untransformed) cell lines. In addition, our experiments unambiguously demonstrate a positive correlation between Galectin-9 levels and invasiveness of cells through ECM. Our results seek to also reconcile the cell-aggregative behavior of Galectin-9 by showing its role in spurring bulk collective (radial) invasion in addition to single cell mesenchymal migration.

The inhibition of invasion phenotypes with Galectin-9 inhibitors with a strong binding affinity for N-CRD (and a relatively weak affinity for C-CRD) implies a relatively more prominent role of the former domain in regulating cancer invasion, but not exclude a role for the latter. Based on exon-intron organization, galectin CRDs have been phylogenetically divided into two types: F4 and F3 CRDs (10). The galectins, whose roles have been well elucidated in a variety of cancers: the prototype Galectin-1 and the chimeric Galectin-3, both possess F3 CRD domain (10). In contrast, the N-CRD of Galectin-9 is an F4 CRD. Along with reports suggesting F4 CRD of Galectin-9 being playing protumorigenic roles (33,34), our study showcases how structurally dissimilar galectin folds may contribute to cancer invasion.

Mesenchymal cancer cells such as MDA-MB-231 adhere to Coll I fibers at their front end and de-adhere at their rear end during migration. The adhesion-deadhesion dynamics is regulated through actin cytoskeletal dynamics which in turn is mediated through the phosphorylation of the Focal Adhesion Kinase (FAK) (35,36). The proinvasive functions of S100A4, proteomically identified to be under the regulation of Galectin-9 levels has been proposed to be associated with its localization and possible interaction with myosin (37) and cytoskeletal elements such as actin (38) and tubulin (39). Our demonstration that Galectin-9 driven accentuation of FAK signaling and its resultant potentiation of S100A4 gene expression, therefore suggests that its effect on the cytoskeletal dynamics of migrating breast epithelia maybe multilevel in nature. Moreover, S100A4 has also been shown to enhance FAK signaling in pancreatic cancer (40), which when taken together with our demonstration of the converse, suggests a positive feedback loop can operate between the two. A possible inductive effect of Galectin-9 on such positive feedback signaling loop may explain its role in promoting cancer cell migration. In this manuscript, we have not explored how Galectin-9 increases FAK phosphorylation. Although its attenuation by the N-CRD binding inhibitors suggests the involvement of the domain, it is unclear whether either of the domains or the linker of Galectin-9 is required to bind FAK directly or indirectly to promote the phosphorylation. We aim to study such intermolecular interactions in the future. Does the C-CRD of Galectin-9 also participate in this function, albeit to a lesser extent than N-CRD? Previous studies show that the two CRDs of Galectin-9 have differential selectivity towards glycoproteins (41). Future investigations will dissect the individual roles of Galectin-9 domains and their linker to shed light on how cancers may adapt the biochemical intricacy of tandem repeat galectins for invasion and metastasis.

## Materials and Methods

### Cell culture

HMLE cells were a kind gift from Dr. Robert Weinberg, Harvard Medical School, and Dr. Annapoorni Rangarajan, Indian Institute of Science. Cells were cultured in DMEM:F12 (1:1) supplemented with 1% fetal bovine serum (Gibco, 10270), 0.5 μg/mL Hydrocortisone (Sigma, H0888), 10 μg/mL Insulin (Sigma, I6634) and 10 ng/mL human recombinant epidermal growth factor (HiMedia, TC228). MCF7 cells were grown in DMEM (HiMedia, AT007F) supplemented with 10% fetal bovine serum. BT-549 & MDA-MB-231 cells were maintained in DMEM:F12 (1:1) (HiMedia, AT140) supplemented with 10% fetal bovine serum (Gibco, 10270). All the cells were cultured in a 37 °C humidified incubator with 5% carbon dioxide.

### RNA isolation

Total RNA was isolated using RNAiso Plus reagent from TaKaRa (TaKaRa, 9108) as mentioned in the product manual. Briefly, the spent medium was aspirated, and cells were washed with 1X PBS. 1 mL of RNAiso Plus reagent is added to the plate, and cells were collected using a cell scraper. Samples were left at room temperature for 5 min. 0.2 mL of chloroform was added, vigorously vortexed, and left at room temperature for 5 min. The upper layer was transferred into a new centrifuge tube following centrifugation at 12,000g for 15 min at 4 °C. 0.5 mL of isopropanol was added and incubated at room temperature for 10 min. After incubation, samples were centrifuged at 12,000g for 10 min at 4 °C, and the supernatant was discarded. The RNA pellet was washed with 1 mL 75% ethanol and centrifuged at 12,000g for 5 min at 4 °C. The supernatant was discarded, and the RNA pellet was air-dried. The RNA pellet was resuspended in appropriate amount of nuclease-free water. RNA samples were stored at -80 °C until further use.

### cDNA synthesis and quantitative real-time PCR

RNA yield was analyzed using the NanoDrop® ND-1000 UV-Vis Spectrophotometer (NanoDrop Technologies, USA). 1 μg of total RNA was reverse transcribed using Verso™ cDNA synthesis kit as per manufacturer’s protocol (Thermo Scientific, AB-1453). Real-time PCR was performed with 1:2 diluted cDNA using SYBR green detection system (Thermo Fischer Scientific, F415L) and Rotorgene Q (Qiagen, 9001560). *18S rRNA* gene was used as internal control for normalization. Relative gene expression was calculated using comparative Ct method. All the genes analyzed along with sequence are mentioned in table 2. Appropriate no template and no-RT control were included in each experiment. All the samples were analyzed in duplicates/triplicates and repeated three times independently.

### Western blotting

Cells were washed twice with ice-cold 1X PBS to remove protein traces from FBS. Cells were lysed using ice-cold radioimmunoprecipitation assay buffer containing protease inhibitor cocktail (Sigma, P8340) and phosphatase inhibitor (Sigma, P5726 & P0044). Cells were collected into a fresh centrifuge tube using a cell scraper. To ensure efficient and complete cell lysis, samples were incubated on ice for 20 min with intermittent vortex. Samples were centrifuged at 13000g for 30 minutes at 4 °C. The clear, cell-debris free supernatant was transferred to a fresh centrifuge tube. Samples were either processed for protein estimation or stored at -80 °C until further use.

Protein concentration was estimated using DC protein assay as mentioned in the manual (Bio-rad Inc., 500-0116). 100 μg of total protein was resolved on 10, 12, or 14% SDS-PAGE gel and transferred onto PVDF membrane (Merck, ISEQ85R) using a semi-dry transfer unit (Bio-Rad Inc, USA) at 0.3 A for 90 min. After transfer, the membrane was blocked with 1X TBST buffer containing 5% BSA protein (HiMedia, MB083) for one hour at room temperature. The membrane was then incubated overnight with primary antibody (for Galectin-9, Ab69630 was used) diluted 1:1000 in 1X TBST containing 5% BSA at 4 °C. The details of primary antibodies used in this study have been mentioned in table 3. The membrane was washed for 10 min in 1X TBST, thrice and incubated with appropriate HRP conjugated secondary antibodies (Sigma, USA) diluted 1:10,000 at room temperature for one hour. The membrane was washed again with 1X TBST for 10 min thrice and developed using chemiluminescent WesternBright® ECL substrate (Advansta, K-12045-C20). The developed blots were imaged and analyzed using ChemiDoc Imaging System (Bio-Rad Inc., USA) at multiple cumulative exposure settings. α-Tubulin expression was used as a loading control in all experiments.

### Invasion assay

8 μm pore-size polycarbonate transwell inserts were obtained from HiMedia (TCP083). Transwells were coated with 200 μg/mL reconstituted basement membrane (rBM or lrECM) (Corning, 354230) as per the manufacturer’s protocol. 3 × 10^4^ cells were seeded in 200 μL of serum-free DMEM:F12 (1:1) medium in each transwell. The bottom well was filled with 1 mL of 10% serum containing DMEM:F12 (1:1) and incubated for 24 h at 37 °C, 5% carbon dioxide containing humidified chamber. Carefully, medium from transwell was removed, washed with 1X PBS once, and cells were fixed using 100% methanol for 10 min at room temperature. After fixing, cells were washed with 1X PBS, and non-invading cells were carefully removed using a moistened cotton swab. Transwells are stained with 1% crystal violet for 15 min at room temperature and washed to remove excess dye. Membranes were dried and imaged under a microscope using 40X total magnification. At least 5 independent fields were imaged per transwell, and number of cells were counted. Each experiment has been performed in duplicates and repeated three times.

### 3D invasion assay

3D invasion assay was performed as described in our earlier work (23). In brief, cancer cells were trypsinized using 1:5 diluted 0.25% Trypsin & 0.02% EDTA (HiMedia, TCL007). 30,000 cells per 200 μL of defined medium (42) supplemented with 4% lrECM (Corning, 354230) were cultured on 3% polyHEMA (Sigma, P3932) coated 96 well plate for 48 h in a 37 °C humidified incubator with 5% carbon dioxide.

lrECM-coated clusters were collected into 1.5 mL tubes, centrifuged briefly, and the supernatant is removed. Acid-extracted rat tail collagen (Gibco, A1048301) was neutralized on ice in the presence of 10X DMEM with 0.1N NaOH such that the final concentration of the collagen is 1 mg/mL. The pellet of clusters was resuspended in 50 μL of neutralized collagen, seeded in 8-well chambered cover glass (Eppendorf 0030742036), and supplemented with a defined medium. 3D cultures were grown in a 37 °C humidified incubator with 5% carbon dioxide.

Endpoint imaging was done after fixing 3D cancer clusters, counterstained with DAPI, Alexa conjugated Phalloidin, and imaged using Carl Zeiss LSM880 confocal microscope with system optimized settings. Brightfield time-lapse imaging of invading cancer clusters was performed on Olympus IX73 fluorescence microscope fitted with stage top incubator and 5% carbon dioxide. Images were collected for 24 h at 10 min intervals.

### Adhesion assay

96 well plates were coated with 50 μg/mL reconstituted basement membrane (rBM or lr-BM) (Corning, 354230) or 50 μg/mL neutralized rat tail collagen (rich in Coll I) (Gibco, A1048301) for 2 hours to overnight at 37 °C. Excess matrix was removed, allowed to dry for 30 min at 37 °C, and blocked with 0.5% BSA (HiMedia, MB083) for 2 h at 37 °C. After blocking, excess BSA was removed, and plates are used for adhesion assay. 0.5% BSA overnight coating at 37 °C was used as a negative control.

Cells were trypsinized & after counting, 30,000 cells per well were incubated in BSA, and ECM coated wells for 30 min at 37 °C. Unadhered cells were removed carefully, and wells were washed with 1X PBS thrice. Cells were fixed using 100% methanol for 10 min at room temperature. After fixing, cells were washed with 1X PBS thrice and stained with 50 μg/mL propidium iodide (HiMedia, TC252) for 30 min at room temperature. Excess stain was removed and washed cells thrice with 1x PBS. Using plate reader, fluorescence was read using Ex 535 nm/Em 617 nm. BSA or ECM without cells was used as blank. The assay was done in triplicates and repeated three times independently.

### Immunohistochemistry

Breast tumor and matched normal sections were made from formalin-fixed paraffin-embedded blocks at Kidwai Cancer Institute, Bangalore, after obtaining necessary approval from the Institutional Human Ethics committee and consent from patients. Sections were incubated at 65 °C overnight to remove wax. Immediately, samples were re-hydrated, gradually incubating in decreasing alcohol concentrations: 2x 5 min Xylene, 2x 5 min 100% Ethanol, 2x 5 min 90% Ethanol, 1x 10min 80% Ethanol, 1x 10 min 70% Ethanol and finally in distilled water for 10 min. Antigen retrieval was performed using citrate buffer pH 6.0 in microwave for 30 min and allowed to cool down to room temperature. Sections were blocked using 5% BSA and 0.01% Tween20 made in 1X PBS pH 7.4 for 1 h at room temperature. Galectin-9 primary antibody (R&D systems, AF-2045) was added to sections at 1 μg/mL dilution and incubated overnight at 4 °C. Sections were washed with 1X PBS + 0.01% Tween20 for 5 min at room temperature thrice. Secondary antibody was added at 1:1000 dilution and incubated at room temperature for 2 h. Sections were washed with 1X PBS + 0.01% Tween20 for 5 min at room temperature thrice. Samples are counterstained with Hematoxylin. Sections were dried using serially incubating in increasing concentrations of alcohol and mounted using DPX.

### Perturbation of *LGALS9* expression

HEK293FT cells were seeded at 50% confluency in DMEM medium supplemented with 10% FBS. After overnight incubation, the medium was replaced with DMEM without FBS for 1 hour before the plasmids pCDH-GAL-9-T2A-Puro (System Biosciences, CD527A-1) or pLKO. 1-shGAL-9 (MISSION® shRNA library, Merck), psPAX2 and pMD2.G (packaging vectors were a kind gift from Dr. Deepak K. Saini at MRDG, Indian Institute of Science) were mixed at a ratio of 2000 ng: 2000 ng: 800 ng, respectively in Turbofect (Invitrogen, USA) before adding to the cell monolayer. PCDH-T2A-Puro (vector control) or pLKO.1-shSc (scrambled control) was used as a control for overexpression and knockdown, respectively. After incubating the cells with the mixture for 6 hours (to allow plasmid uptake), the media was replaced with DMEM with 10% FBS. Medium containing lentiviral particles was then harvested at 48 hours and again at 72 hours after transfection and used directly for viral transduction or stored at -80oC.

To obtain MDA-MB-231 cells stably overexpressing or knockdown of Galectin-9, cells were seeded at 50% confluency. After overnight incubation, cells were treated with DMEM:F12 (1:1)+10% FBS containing Polybrene (Sigma, USA) at a final concentration of 4 μg/mL. Medium containing lentiviral particles was then added to the cell monolayer and incubated for 48 hours. At the end of incubation, transduced cells were selected by incubating the monolayer in DMEM:F12 (1:1)+10% FBS containing 2 μg/mL Puromycin dihydrochloride (Sigma, USA) for 24 h. The same Puromycin pressure was maintained for two passages and then increased to 5 μg/mL for another two passages before being withdrawn. Stably transduced cells were confirmed for expression using western blot or quantitative real-time PCR.

### Synthesis procedures, and data for the synthesis of 2 and 3

#### General Procedures

All glassware was oven dried. All solvents and reagents were purchased from commercial sources or synthesized via literature protocols and used without further purification. TLC analysis was performed on precoated Merck silica gel 60 F_254_ plates using UV light and charring solution (10 mL conc. H_2_SO_4_/90 mL EtOH). Flash column chromatography was performed on SiO_2_ purchased from Aldrich (technical grade, 60 Å pore size, 230–400 mesh, 40–63 μm). All microwave-assisted organic synthesis was performed in a Biotage® Initiator+ (0–400 W from magnetron at 2.45 GHz), or in a Biotage® Initiator Classic (0–400 W from magnetron at 2.45 GHz). Final compounds were purified using preparative HPLC on an Agilent 1260 Infinity system with a Symmetry Prep C_18_, 5 μM, 19 mm × 100 mm column using a gradient (water with 0.1% formic acid and acetonitrile), 0-5 min with 0-10% acetonitrile, 5-10 min with 10-60% acetonitrile, 10-13 min with 60-90% acetonitrile, 13-15 min with 90-100% acetonitrile, and 15-20 min with 100% acetonitrile. Monitoring and collection were based on UV-vis absorbance at 210 and 254 nm. All NMR spectra were recorded with a Bruker DRX 400 MHz spectrometer (400 MHz for ^1^H, 101 MHz for ^13^C) and Varian 500 MHz (500 MHz for ^1^H, 126 MHz for ^13^C) at ambient temperature using CD_3_OD as the solvent. Chemical shifts are given in ppm relative to the residual solvent peak (^1^H NMR: CD_3_OD δ 3.31; ^13^C NMR: CD_3_OD δ 49.00) with multiplicity (b = broad, s = singlet, d = doublet, t = triplet, q = quartet, quin = quintet, hept = heptet, m = multiplet, app = apparent), coupling constants (in Hz) and integration. High-resolution mass analyses were performed using a Micromass Q-TOF mass spectrometer (ESI).

#### 3,4-Dichlorophenyl 3-*O*-[3-(3,4,5-trifluorobenzylamino)-3-(4-methylbenzensulfonylimino)-propyl] β-D-galactopyranoside (2)

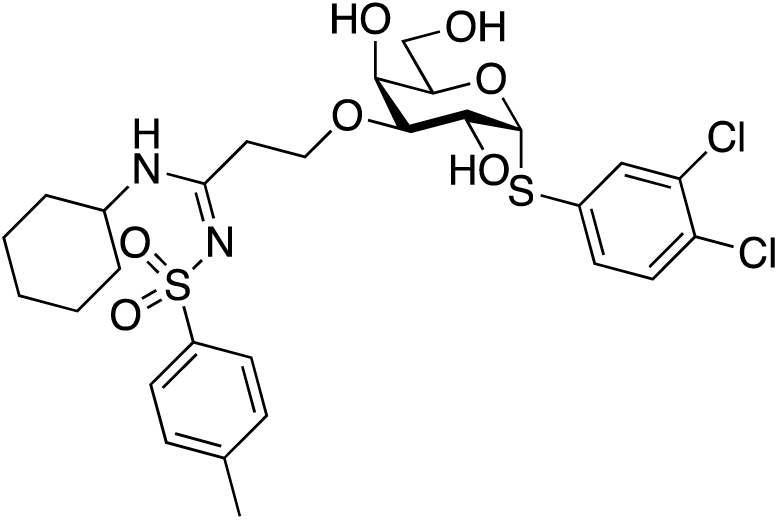

3,4-Dichlorophenyl 3-*O*-propargyl-1-thio-α-D-galactopyranoside **1** (26) (20.4 mg, 0.054 mmol) was dissolved in dry THF (1.0 mL). Cyclohexylamine (31.1 μL, 273 μmol) and 4-methylbenzenesulfonyl azide (43.2 μL, 198 μmol) were added and the mixture was degassed by purging with N_2_. CuI (10.7 mg, 0.056 mmol) was added to a separate round bottom flask, the flask was flushed with N_2_ gas, dry THF (1.0 mL) was added, and the mixture was sonicated. The solution containing CuI was transferred to the solution containing **1**, cyclohexylamine, and 4-methylbenzenesulfonyl azide. The clear transparent reaction mixture was stirred for 1.5 h, then concentrated in vacuo. Flash column chromatography of the residue (EtOAc:n-Heptane 5:1) gave **3** (25.7.3 mg, 74%), which was further purified by preparative HPLC. ^1^H NMR (CD_3_OD, 400 MHz): δ 7.75 (d, *J* = 8.3 Hz, 2H, Ph), 7.73 (d, *J* = 2.0 Hz, 1H, Ph), 7.48 (dd, *J* = 8.6, 2.0 Hz, 1H, Ph), 7.45 (d, *J* = 8.3 Hz, 1H, Ph), 7.35 (d, *J* = 8.0 Hz, 2H, Ph), 5.66 (d, *J* = 5.6 Hz, 1H, H-1), 4.31-4.23 (m, 2H, H-2 and H-5), 4.14 (d, *J* = 3.0 Hz, 1H, H-4), 3.93 (m, 1H, CH_2_), 3.82 (m, 1H, CH), 3.79-3.69 (m, 3H, H-6, H-6’ and CH_2_), 3.43 (dd, *J* = 10.2, 3.1 Hz, 1H, H-3), 3.01 (m, 2H, CH_2_), 2.42 (s, 3H, CH_3_), 1.91 (m, 2H, CH_2_), 1.75 (m, 2H, CH_2_), 1.64 (m, 2H, CH_2_), 1.40-1.13 (m, 4H, CH_2_). ^13^C NMR (CD_3_OD, 100 MHz): δ 167.3, 143.8, 142.5, 136.9, 133.5, 132.7, 132.1, 131.7, 130.4, 130.4, 127.1, 91.1, 80.3, 73.4, 68.7, 67.2, 66.7, 62.4, 52.3, 34.8, 32.8, 32.8, 26.5, 26.0, 21.4 HRMS calculated for [C_28_H_37_Cl_2_N_2_O_7_S_2_]^+^, 647.1419; found: 647.1424. HPLC purity by UV/VIS detector (254 nm): 95.0%

#### 3,4-Dichlorophenyl 3-*O*-[3-(4-fluorobenzylamino)-3-(4-methylbenzensulfonylimino)-propyl]-β-D galactopyranoside (3)

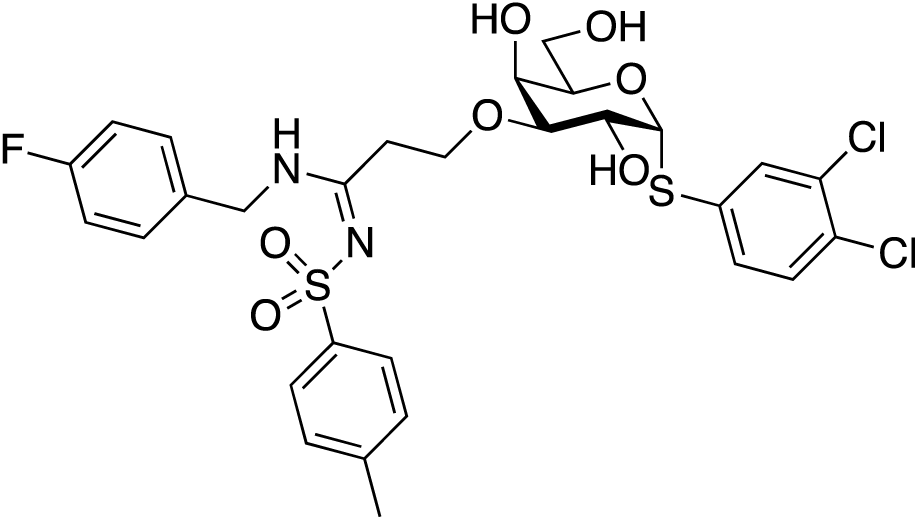

3,4-Dichlorophenyl 3-*O*-propargyl-1-thio-α-D-galactopyranoside **1** (26) (73.9 mg, 0.195 mmol) was dissolved in dry THF (10 mL). 4-Fluorobenzylamine (67 μL, 586 μmol) and 4-methylbenzenesulfonyl azide (128 μL, 584 μmol) were added and the mixture was degassed by purging with N_2_. CuI (38.8 mg, 0.204 mmol) was added to a separate round bottom flask, the flask was flushed with N2 gas, dry THF (10 mL) was added, and the mixture was sonicated. DIPEA (0.679 ml, 3.898 mmol) was added and the solution containing CuI and DIPEA was transferred to the solution containing **1**, 4-fluorobenzylamine, and 4-methylbenzenesulfonyl azide. The transparent yellow reaction mixture was stirred at 50°C overnight, then concentrated in vacuo. Flash column chromatography of the residue (EtOAc:n-Heptane 3:1, followed by EtOAc:n-acetone 1:2) gave **3** (76.3 mg, 58%), which was further purified by preparative HPLC. ^1^H NMR (500 MHz, MeOD) δ 7.73 (d, J = 2.0 Hz, 1H, Ph), 7.67 – 7.59 (m, 2H, Ph), 7.50 – 7.41 (m, 2H, Ph), 7.34 – 7.26 (m, 4H, Ph), 7.05 – 6.95 (m, 2H, Ph), 5.66 (d, J = 5.5 Hz, 1H, H-1), 4.49 – 4.39 (m, 2H, CH_2_),, 4.31 – 4.23 (m, 2H, H-2 and H-5), 4.14 (dd, *J* = 3.2, 1.3 Hz, 1H, H-4), 3.99 (ddd, *J* = 10.1, 7.0, 4.5 Hz, 1H, CH_2_), 3.79 (ddd, *J* = 10.0, 6.6, 4.6 Hz, 1H, CH_2_), 3.76 – 3.65 (m, 2H, H-6, H-6’), 3.46 (dd, *J* = 10.1, 3.1 Hz, 1H, H-3), 3.09 (td, *J* = 6.9, 4.5 Hz, 2H, CH_2_), 2.41 (s, 3H, CH_3_). ^13^C NMR (126 MHz, MeOD) δ 167.3, 166.9, 143.7, 142.9, 142.7, 137.0, 134.4, 134.4, 133.5, 132.7, 132.7, 132.1, 131.7, 130.4, 130.4, 127.1, 127.0, 91.3, 91.2, 80.7, 80.6, 73.6, 73.6, 68.9, 68.8, 67.4, 67.3, 67.2, 67.0, 62.4, 59.5, 58.4, 49.6, 33.2, 32.5, 32.0, 31.8, 31.7, 30.3, 30.3, 26.7, 26.5, 26.4, 26.4, 26.3, 21.4. HRMS: m/z calcd for (M + Na)+ 673.1012, found 673.1027. HPLC purity by UV/VIS detector (254 nm): 95.5%

### Galectin-9 inhibition

The compounds **2** and **3** are dissolved in DMSO and diluted in complete medium for experiments. Through the study, MDA-MB-231 cells were treated with Galectin-9 inhibitors **2** and **3** at a viability sub-IC50 concentration of 10 μM obtained from the dose-response curves (IC50 of 18 and 19 μM for **2** and **3**, respectively). For adhesion assay, cells were treated with **2** and **3** for one hour. For 3D invasion assay, cells were incubated with the inhibitors throughout the 24 h time. Appropriate vehicle (DMSO) control was used for the experiments.

### FAK inhibition

FAK was inhibited using 1,2,4,5-Benzenetetramine tetrahydrochloride commonly known as FAK inhibitor 14 (4506-66-5, Sigma, 14485). MDA-MB-231 cells overexpressing Galectin-9 was either treated with vehicle control (DMSO) or 100 nM of the inhibitor for 6 hour and S100A4 gene expression was analysed using qRT-PCR.

### Mass spectrometry

25 μL samples were taken and reduced with 5 mM tris(2-carboxyethyl) phosphine (TCEP) and further alkylated with 50 mM iodoacetamide and then digested with Trypsin (1:50, trypsin/lysate ratio) for 16 h at 37 °C. Digests were cleaned using a C18 silica cartridge to remove the salt and dried using a speed vac. The dried pellet was resuspended in buffer A (5% acetonitrile, 0.1% formic acid). All the experiment was performed using EASY-nLC 1,000 system (Thermo Fisher Scientific) coupled to Thermo Fisher-QExactive equipped with nanoelectrospray ion source. 1.0 μg of the peptide mixture was resolved using 15 cm PicoFrit column (360 μm outer diameter, 75 μm inner diameter, 10 μm tip) filled with 2.0 μm of C18-resin (Dr Maeisch). The peptides were loaded with buffer A and eluted with a 0–40% gradient of buffer B (95% acetonitrile, 0.1% formic acid) at a flow rate of 300 nl/min for 100 min. MS data were acquired using a data-dependent top10 method dynamically choosing the most abundant precursor ions from the survey scan. All samples were processed, and RAW files generated were analyzed with Proteome Discoverer (v2.2) against the Uniprot HUMAN reference proteome database. For Sequest search, the precursor and fragment mass tolerances were set at 10 ppm and 0.5 D, respectively. The protease used to generate peptides, that is, enzyme specificity was set for trypsin/P (cleavage at the C terminus of “K/R”: unless followed by “P”) along with maximum missed cleavages value of two. Carbamidomethyl on cysteine as fixed modification and oxidation of methionine and N-terminal acetylation were considered as variable modifications for database search. Both peptide spectrum match and protein false discovery rate were set to 0.01 FDR. Statistical analysis was performed by using in-house R script. Abundance value for each run (including all biological replicates) were filtered and imputed by using normal distribution. Log2 transformed abundance values were normalized using Z-score. ANOVA and t-test was performed based on P-value (threshold P < 0.05) to identify the significant proteins.

### Image analysis

#### a. Collective cell invasion

Collective cell invasion was analyzed using ImageJ (43). Image at 0 hour was selected, and cluster area was obtained after auto threshold. This is termed as initial area (A). Similarly, cluster area was obtained at the endpoint, and the area is termed the final area (B). Area of collective cell invasion is calculated using the formula: (B-A)/(A).

#### b. Single-cell invasion

Image at endpoint was obtained, and the number of single cells dispersed in Coll I was counted manually using cell counter plugin in ImageJ.

#### c. Velocity and accumulated distance

Velocity and accumulated distance were calculated using the manual tracking plugin. Briefly, single-cells coming out of the cluster was tracked manually for the last 12 hour of the time-lapse videography. Tracks obtained from such single cells were analyzed using the Chemotaxis tool by ibidi, Germany.

### Statistical analysis

All experiments were performed in at least duplicates and repeated thrice independently. All data is represented as mean±SEM unless specified. For statistical analysis, unpaired student’s *t*-test with Welch’s correction or One-way ANOVA with Dunnett’s multiple comparisons was performed. Statistical significance is represented using P-value: * P≤0.05, **P≤0.01, ***P≤0.001, ****P<0.0001.

## Acknowledgments

D.P. is supported by Senior Research Fellowship (SRF) from Ministry of Education, India. This work was supported by the Wellcome Trust/DBT India Alliance Fellowship/Grant IA/I/17/2/ 503312] awarded to R.B. In addition, this work was supported by funds from the Department of Biotechnology, India [BT/ 909 PR26526/GET/119/92/2017], SERB[ECR/2015/000280], DBT-IISc partnership program (BT/PR27952/INF/22/212/2018), and the Institute of Eminence grant (IE/CARE-19-0319). UN was supported by funds from The Swedish Research Council (Grant No. 621-2016-03667). UN and HL were supported by funds from Galecto Biotech Inc. The authors thank Barbro Kahl Knutson for the fluorescence polarization measurements and Sofia Essén for high-resolution mass spectrometry and purity determinations. We thank the Bioimaging facility at Division of Biological Science, Indian Institute of Science for help with confocal microscopy.

## Author Contribution

DP and RB designed the biological experiments. DP, MB, and SH performed the biological experiments. DP, MB, SH, RVK, AP, ER, LG, KP, UJN, HL and RB analyzed the data and wrote the manuscript. UJN conceptualized the inhibitor development, did the inhibitor design, supervised the synthesis, analysed galectin binding evaluations, and wrote the inhibitor design and synthesis and galectin binding analysis parts. HL conceptualized the galectin binding assay and analysed data from it. AP, ER, LG, and KP, were involved in design and synthesis of the inhibitors, wrote the synthesis experimental part, and analysed galectin binding data.

## References

1. Barondes, S. H., Cooper, D. N., Gitt, M. A., and Leffler, H. (1994) Galectins. Structure and function of a large family of animal lectins. The Journal of biological chemistry 269, 20807–20810

2. Cummings, R. D., Liu, F. T., and Vasta, G. R. (2015) Galectins. in Essentials of Glycobiology (rd, Varki, A., Cummings, R. D., Esko, J. D., Stanley, P., Hart, G. W., Aebi, M., Darvill, A. G., Kinoshita, T., Packer, N. H., Prestegard, J. H., Schnaar, R. L., and Seeberger, P. H. eds.), Cold Spring Harbor (NY). pp 469-480

3. Liu, F. T., and Rabinovich, G. A. (2005) Galectins as modulators of tumour progression. Nature reviews. Cancer 5, 29–41

4. Johannes, L., Jacob, R., and Leffler, H. (2018) Galectins at a glance. Journal of cell science 131

5. Di Lella, S., Sundblad, V., Cerliani, J. P., Guardia, C. M., Estrin, D. A., Vasta, G. R., and Rabinovich, G. A. (2011) When galectins recognize glycans: from biochemistry to physiology and back again. Biochemistry 50, 7842–7857

6. Arthur, C. M., Baruffi, M. D., Cummings, R. D., and Stowell, S. R. (2015) Evolving mechanistic insights into galectin functions. Methods in molecular biology 1207, 1–35

7. Moar, P., and Tandon, R. (2021) Galectin-9 as a biomarker of disease severity. Cellular immunology 361, 104287

8. Thijssen, V. L., Heusschen, R., Caers, J., and Griffioen, A. W. (2015) Galectin expression in cancer diagnosis and prognosis: A systematic review. Biochimica et biophysica acta 1855, 235–247

9. Cooper, D. N., and Barondes, S. H. (1999) God must love galectins; he made so many of them. Glycobiology 9, 979–984

10. Houzelstein, D., Goncalves, I. R., Fadden, A. J., Sidhu, S. S., Cooper, D. N., Drickamer, K., Leffler, H., and Poirier, F. (2004) Phylogenetic analysis of the vertebrate galectin family. Molecular biology and evolution 21, 1177–1187

11. Dai, S. Y., Nakagawa, R., Itoh, A., Murakami, H., Kashio, Y., Abe, H., Katoh, S., Kontani, K., Kihara, M., Zhang, S. L., Hata, T., Nakamura, T., Yamauchi, A., and Hirashima, M. (2005) Galectin-9 induces maturation of human monocyte-derived dendritic cells. Journal of immunology 175, 2974–2981

12. Kashio, Y., Nakamura, K., Abedin, M. J., Seki, M., Nishi, N., Yoshida, N., Nakamura, T., and Hirashima, M. (2003) Galectin-9 induces apoptosis through the calcium-calpain-caspase-1 pathway. Journal of immunology 170, 3631–3636

13. Smith, L. K., Fawaz, K., and Treanor, B. (2021) Galectin-9 regulates the threshold of B cell activation and autoimmunity. eLife 10

14. Selno, A. T. H., Schlichtner, S., Yasinska, I. M., Sakhnevych, S. S., Fiedler, W., Wellbrock, J., Klenova, E., Pavlova, L., Gibbs, B. F., Degen, M., Schnyder, I., Aliu, N., Berger, S. M., Fasler-Kan, E., and Sumbayev, V. V. (2020) Transforming growth factor beta type 1 (TGF-beta) and hypoxia-inducible factor 1 (HIF-1) transcription complex as master regulators of the immunosuppressive protein galectin-9 expression in human cancer and embryonic cells. Aging 12, 23478–23496

15. Grosset, A. A., Labrie, M., Vladoiu, M. C., Yousef, E. M., Gaboury, L., and St-Pierre, Y. (2016) Galectin signatures contribute to the heterogeneity of breast cancer and provide new prognostic information and therapeutic targets. Oncotarget 7, 18183–18203

16. Irie, A., Yamauchi, A., Kontani, K., Kihara, M., Liu, D., Shirato, Y., Seki, M., Nishi, N., Nakamura, T., Yokomise, H., and Hirashima, M. (2005) Galectin-9 as a prognostic factor with antimetastatic potential in breast cancer. Clinical cancer research : an official journal of the American Association for Cancer Research 11, 2962–2968

17. Zhou, X., Sun, L., Jing, D., Xu, G., Zhang, J., Lin, L., Zhao, J., Yao, Z., and Lin, H. (2018) Galectin-9 Expression Predicts Favorable Clinical Outcome in Solid Tumors: A Systematic Review and Meta-Analysis. Frontiers in physiology 9, 452

18. Chandrashekar, D. S., Bashel, B., Balasubramanya, S. A. H., Creighton, C. J., Ponce-Rodriguez, I., Chakravarthi, B., and Varambally, S. (2017) UALCAN: A Portal for Facilitating Tumor Subgroup Gene Expression and Survival Analyses. Neoplasia 19, 649–658

19. Gadwal, A., Modi, A., Khokhar, M., Vishnoi, J. R., Choudhary, R., Elhence, P., Banerjee, M., and Purohit, P. (2021) Critical appraisal of epigenetic regulation of galectins in cancer. International journal of clinical oncology

20. Pally, D., Pramanik, D., Hussain, S., Verma, S., Srinivas, A., Kumar, R. V., Everest-Dass, A., and Bhat, R. (2021) Heterogeneity in 2,6-Linked Sialic Acids Potentiates Invasion of Breast Cancer Epithelia. ACS central science 7, 110–125

21. Wolf, K., Mazo, I., Leung, H., Engelke, K., von Andrian, U. H., Deryugina, E. I., Strongin, A. Y., Brocker, E. B., and Friedl, P. (2003) Compensation mechanism in tumor cell migration: mesenchymal-amoeboid transition after blocking of pericellular proteolysis. The Journal of cell biology 160, 267–277

22. Friedl, P., Locker, J., Sahai, E., and Segall, J. E. (2012) Classifying collective cancer cell invasion. Nature cell biology 14, 777–783

23. Pally, D., Pramanik, D., and Bhat, R. (2019) An Interplay Between Reaction-Diffusion and Cell-Matrix Adhesion Regulates Multiscale Invasion in Early Breast Carcinomatosis. Frontiers in physiology 10, 790

24. Zetterberg, F. R., Peterson, K., Johnsson, R. E., Brimert, T., Håkansson, M., Logan, D. T., Leffler, H., and Nilsson, U. J. (2018) Monosaccharide Derivatives with Low-Nanomolar Lectin Affinity and High Selectivity Based on Combined Fluorine-Amide, Phenyl-Arginine, Sulfur-π, and Halogen Bond Interactions. ChemMedChem 27, 133–137

25. Delaine, T., Collins, P., MacKinnon, A., Sharma, G., Stegmayr, J., Rajput, V. K., Mandal, S., Cumpstey, I., Larumbe, A., Salameh, B. A., Kahl-Knutsson, B., van Hattum, H., van Scherpenzeel, M., Pieters, R. J., Sethi, T., Schambye, H., Oredsson, S., Leffler, H., Blanchard, H., and Nilsson, U. J. (2016) Galectin-3-Binding Glycomimetics that Strongly Reduce Bleomycin-Induced Lung Fibrosis and Modulate Intracellular Glycan Recognition. Chembiochem : a European journal of chemical biology 17, 1759–1770

26. Dahlqvist, A., Zetterberg, F. R., Leffler, H., and Nilsson, U. J. (2019) Aminopyrimidine-galactose hybrids are highly selective galectin-3 inhibitors. MedChemComm 10, 913–925

27. Sorme, P., Kahl-Knutsson, B., Huflejt, M., Nilsson, U. J., and Leffler, H. (2004) Fluorescence polarization as an analytical tool to evaluate galectin-ligand interactions. Anal Biochem 334, 36–47

28. Pylayeva, Y., Gillen, K. M., Gerald, W., Beggs, H. E., Reichardt, L. F., and Giancotti, F. G. (2009) Ras- and PI3K-dependent breast tumorigenesis in mice and humans requires focal adhesion kinase signaling. The Journal of clinical investigation 119, 252–266

29. Krakhmal, N. V., Zavyalova, M. V., Denisov, E. V., Vtorushin, S. V., and Perelmuter, V. M. (2015) Cancer Invasion: Patterns and Mechanisms. Acta naturae 7, 17–28

30. Sieg, D. J., Hauck, C. R., and Schlaepfer, D. D. (1999) Required role of focal adhesion kinase (FAK) for integrin-stimulated cell migration. Journal of cell science 112 (Pt 16), 2677–2691

31. Che, P., Yang, Y., Han, X., Hu, M., Sellers, J. C., Londono-Joshi, A. I., Cai, G. Q., Buchsbaum, D. J., Christein, J. D., Tang, Q., Chen, D., Li, Q., Grizzle, W. E., Lu, Y. Y., and Ding, Q. (2015) S100A4 promotes pancreatic cancer progression through a dual signaling pathway mediated by Src and focal adhesion kinase. Scientific reports 5, 8453

32. Zhao, X. K., Cheng, Y., Liang Cheng, M., Yu, L., Mu, M., Li, H., Liu, Y., Zhang, B., Yao, Y., Guo, H., Wang, R., and Zhang, Q. (2016) Focal Adhesion Kinase Regulates Fibroblast Migration via Integrin beta-1 and Plays a Central Role in Fibrosis. Scientific reports 6, 19276

33. Grosset, A. A., Poirier, F., Gaboury, L., and St-Pierre, Y. (2016) Galectin-7 Expression Potentiates HER-2-Positive Phenotype in Breast Cancer. PloS one 11, e0166731

34. Trebo, A., Ditsch, N., Kuhn, C., Heidegger, H. H., Zeder-Goess, C., Kolben, T., Czogalla, B., Schmoeckel, E., Mahner, S., Jeschke, U., and Hester, A. (2020) High Galectin-7 and Low Galectin-8 Expression and the Combination of both are Negative Prognosticators for Breast Cancer Patients. Cancers 12

35. Odenthal, J., Takes, R., and Friedl, P. (2016) Plasticity of tumor cell invasion: governance by growth factors and cytokines. Carcinogenesis 37, 1117–1128

36. Friedl, P., and Alexander, S. (2011) Cancer invasion and the microenvironment: plasticity and reciprocity. Cell 147, 992–1009

37. Kim, E. J., and Helfman, D. M. (2003) Characterization of the metastasis-associated protein, S100A4. Roles of calcium binding and dimerization in cellular localization and interaction with myosin. The Journal of biological chemistry 278, 30063–30073

38. Watanabe, Y., Usada, N., Minami, H., Morita, T., Tsugane, S., Ishikawa, R., Kohama, K., Tomida, Y., and Hidaka, H. (1993) Calvasculin, as a factor affecting the microfilament assemblies in rat fibroblasts transfected by src gene. FEBS letters 324, 51–55

39. Lakshmi, M. S., Parker, C., and Sherbet, G. V. (1993) Metastasis associated MTS1 and NM23 genes affect tubulin polymerisation in B16 melanomas: a possible mechanism of their regulation of metastatic behaviour of tumours. Anticancer research 13, 299–303

40. Kuper, C., Beck, F. X., and Neuhofer, W. (2014) NFAT5-mediated expression of S100A4 contributes to proliferation and migration of renal carcinoma cells. Frontiers in physiology 5, 293

41. Cederfur, C., Salomonsson, E., Nilsson, J., Halim, A., Oberg, C. T., Larson, G., Nilsson, U. J., and Leffler, H. (2008) Different affinity of galectins for human serum glycoproteins: galectin-3 binds many protease inhibitors and acute phase proteins. Glycobiology 18, 384–394

42. Blaschke, R. J., Howlett, A. R., Desprez, P. Y., Petersen, O. W., and Bissell, M. J. (1994) Cell differentiation by extracellular matrix components. Methods in enzymology 245, 535–556

43. Rueden, C. T., Schindelin, J., Hiner, M. C., DeZonia, B. E., Walter, A. E., Arena, E. T., and Eliceiri, K. W. (2017) ImageJ2: ImageJ for the next generation of scientific image data. BMC bioinformatics 18, 529

