## Supplementary file for "Galectin-9 signaling drives breast cancer invasion through matrix"

**Figure S1:**

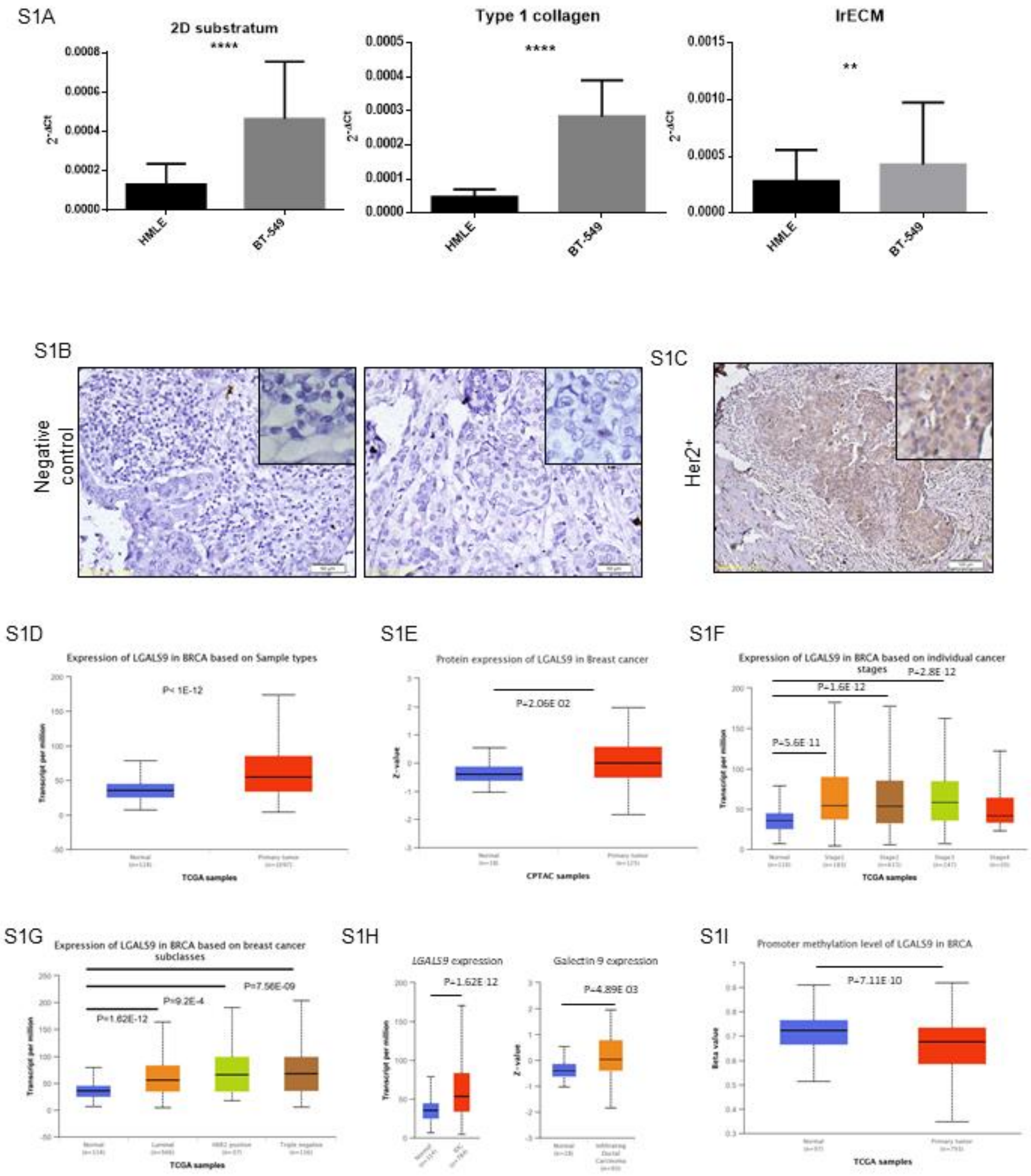


Figure S1: A) Bar graph showing significant increase in Galectin-9 gene expression in BT-549 compared to HMLE when cells were cultured as monolayer (left), non-fibrillar lrECM (middle) and fibrillar collagen I (right). n=3, Mean±SEM. B) Representative micrographs of primary antibody omitted (negative control) breast cancer sections showing no staining. Sections are counterstained with Haematoxylin (blue). C) Immunohistochemical staining showing Galectin-9 (brown) levels in HER2+ breast cancer section. Section is counterstained with Haematoxylin (blue). D) Graph showing significantly higher Galectin-9 mRNA expression in primary breast cancer samples compared to normal samples analyzed using RNAseq. E) Graph showing higher Galectin-9 protein levels in primary tumor samples compared to normal samples analyzed using immunohistochemistry. F) Graph depicting higher Galectin-9 mRNA levels across all stages during breast cancer progression compared to normal samples. G) Graph showing significantly higher Galectin-9 transcript levels in Luminal, Her2+ and triple negative subtypes of breast cancer than normal samples. H) Graphs showing higher Galectin-9 mRNA (left) and protein (right) levels in infiltrating ductal carcinoma samples (IDC) compared to normal samples. I) Bar graph showing lower promoter methylation in primary tumor samples than normal samples correlating with higher Galectin-9 expression in primary tumor (D-I: represents data from level 3 TCGA RNA-seq using ualcan.path.uab.edu. Data is represented as box and whisker plots showing minimum, 25th percentile, median, 75th percentile and maximum values. Outliers are excluded from the plots. *t*-test was performed to ascertain statistical significance).

**Figure S2:**

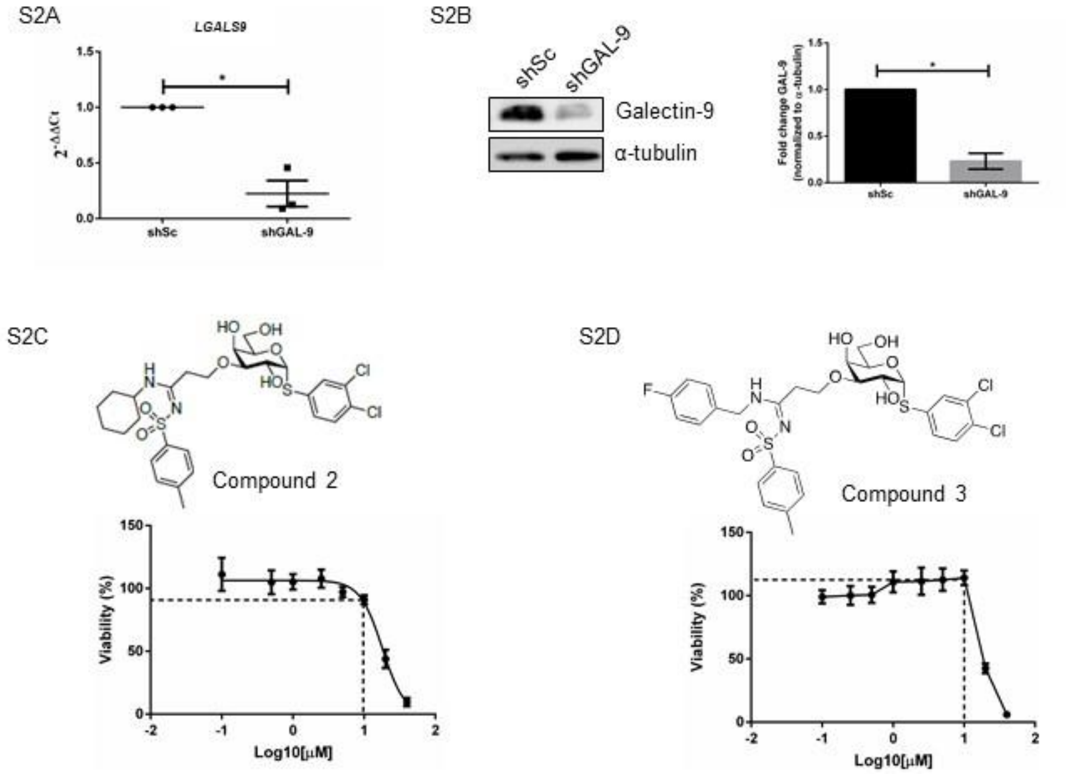


Figure S2: A) Graph showing significantly lower Galectin-9 transcript levels in Galectin-9 depleted (shGAL-9) MDA-MB-231 cells compared to scrambled control cells. B) Immunoblot (left) showing lower Galectin-9 protein levels in shGAL-9 cells compared to scrambled control cells. Bar graph (right) showing quantification of Galectin-9 levels from immunoblot normalized to α-tubulin. n=3, Mean±SEM. Unpaired Student's *t*-test with Welch's correction. * P≤0.05. C) Chemical structure of **3** (top) and graph depicting viability of MDA-MB-231 cells (bottom) at various concentrations ranging from 0.01 to 40 µM at 24 h. Dashed lines indicate the concentration (vertical) used in this study and respective viability of cells (horizontal). D) Chemical structure of **4** (top) and graph depicting viability of MDA-MB-231 cells (bottom) at various concentrations ranging from 0.01 to 40 µM at 24 h. Dashed lines indicate the concentration (vertical) used in this study and respective viability of cells (horizontal).

**Figure S3:**

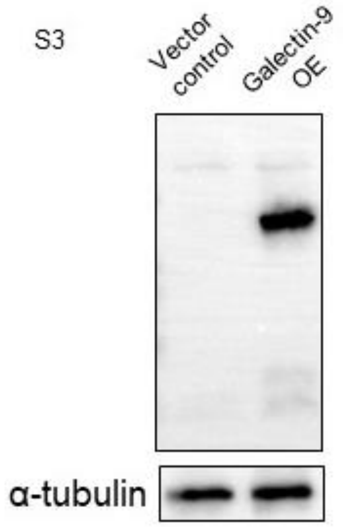


Figure S3: Immunoblot showing higher Galectin-9 expression after overexpression the cDNA in MDA-MB-231 cells.

**Figure S4:**

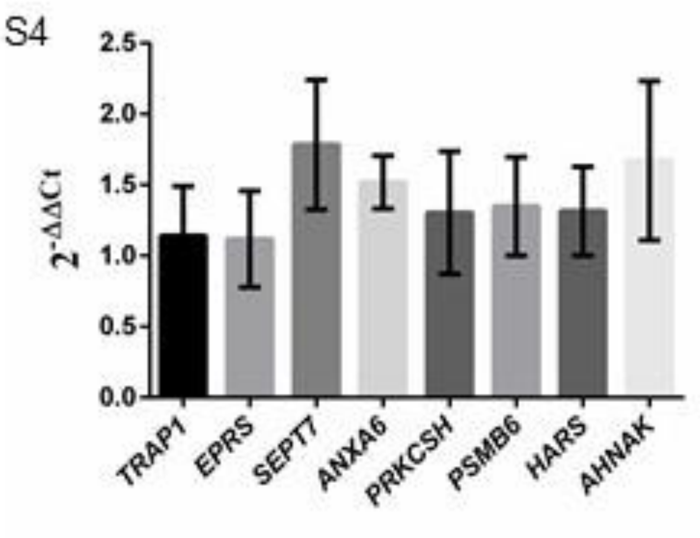


Figure S4: Bar graph showing no change in transcript levels of various genes that are shown to be up-regulated using quantitative proteomics in Galectin-9 overexpressing cells. n=2, Mean±SEM.

**Table 2:**

| Gene | Forward (5'-3') | Reverse (5'-3') |
|---|---|---|
| *LGALS9* | GAGTCCAGCTGTCCCCTTTT | TTGCACACCACGTACCCTC |
| *18S rRNA* | GTAACCCGTTGAACCCCATT | CCATCCAATCGGTAGTAGCG |
| *EPRS* | CTCCTGCTGAACAGATGAAAGC | TTTGCAGCGATAAAGGGTTGG |
| *SEPT7* | GAAGCGTTTGCATGAAAAAGTGA | AACTGTTGGCATTCCTCTGGT |
| *PSMB6* | ACTGACAAGCTGACACCTATTCA | CGGTATCGGTAACACATCTCCTT |
| *ANXA6* | CTGGTGGCAGCATACAAAGAT | CGCTCACTACGTCATCCTCC |
| *PRKCSH* | GGATGGGGCGTTGTCAGAAG | GACGCGGTCGTAGAAAGAGG |
| *TRAP1* | AGGTGTTTATACGGGAGCTGA | GCATTGGTCTGCAAGTGAATCTC |
| *HARS* | ACGTAATCATCCGTTGCTTCAA | ATCCCGCCGATATACCTTTGC |
| *S100A4* | GATGAGCAACTTGGACAGCAA | CTGGGCTGCTTATCTGGGAAG |
| *AHNAK* | GGTAGGCCAGTAGAGGTACAG | CCCCACAGAGACTTCAGGT |

Table 2: List of primers used in the study.

**Table 3:**

| Antibody | Make | Catalog no. | Dilution |
|---|---|---|---|
| Anti-Galectin-9 (Western blotting) | Abcam (UK) | Ab69630 | 1:1000 |
| Anti-Galectin-9 (Immunohistochemistry) | R&D systems | AF2045 | 1 µg/mL |
| Anti- α-tubulin | MERCK | CP06 | 1:5000 |
| Phospho-FAK (Tyr397) | Cell Signaling Technology, Inc. (USA) | 8556 | 1:1000 |
| Total FAK | Cell Signaling Technology, Inc. (USA) | 3285 | 1:1000 |

Table 3: List of antibodies used in the study.
